# Extracellular Vesicle Carryover Distorts Nanoparticle Protein Corona Profiles in Human Plasma

**DOI:** 10.64898/2026.02.19.706828

**Authors:** Bahareh Ghaffari, Shaun Grumelot, Seyed Amirhossein Sadeghi, Alphan Alpaydin, Kylie Hilsen, Brooke Shango, Danilo Ritz, Alexander Schmidt, Hojatollah Vali, Liangliang Sun, Catherine J. Murphy, Amir Ata Saei, Babak Borhan, Morteza Mahmoudi

## Abstract

The nanoparticle (NP) protein corona is considered the biological identity that determines NP fate, safety, targeting, and therapeutic effectiveness in biofluids. Nonetheless, standard corona isolation workflows assume the recovered protein signature originates primarily from plasma proteins adsorbed directly onto the NP surface, while largely overlooking co-isolation of endogenous nanoscale biological structures such as extracellular vesicles (EVs). This oversight can distort the apparent “biological identity” of the NP. Here, we show that EVs are a major hidden contributor to the perceived protein corona composition in human plasma. Using highly monodispersed polystyrene NPs (50-1000 nm) and superparamagnetic beads, we compared corona formation in standard human plasma and plasma depleted of an EV-enriched sedimentable fraction by ultracentrifugation at 100,000 × g for 2 h, with the recovered vesicles subsequently characterized by MACSPlex immunoaffinity analysis. Mass spectrometry revealed that EV depletion reduced the number of proteins identified on polystyrene NPs by 60-75% and on magnetic beads by 45-50%, demonstrating a substantial fraction of the conventionally assigned corona proteome arises from EV-associated carryover. EV depletion also restructured the apparent abundance hierarchy, increasing the relative prominence of soluble plasma proteins such as albumin and shifting dominant signals away from intracellular cytoskeletal component proteins that are characteristic of EV carryover towards genuine soluble plasma proteins and complement factors. These results highlight that standard corona workflows can inadvertently co-isolate a vast array of EV-associated material and thereby yield inaccurate assignments of protein origin. Distinguishing proteins adsorbed from the soluble phase from EV-surface and intravesicular material is essential for accurate interpretation of NP-biofluid interactions, biomarker discovery, and therapeutic targeting because molecular compartment determines both analytical meaning and drug accessibility.

**Significance Statement:** The nanoparticle (NP) “protein corona” defines how engineered nanomaterials interact with living systems, influencing therapeutic safety, efficacy, and diagnostic utility. Conventionally, corona isolation workflows assume that recovered proteins were adsorbed directly from the fluid phase. This study reveals a major, previously overlooked source of analytical distortion: standard separation techniques routinely co-isolate endogenous extracellular vesicles (EVs), drastically distorting the perceived biological identity of NPs. Depletion of an EV-enriched sedimentable fraction by ultracentrifugation reduced identified corona proteins by up to 75% and restructures the apparent proteomic hierarchy. Distinguishing true soluble adsorbates from vesicular carryover is essential for accurately predicting NP behavior in vivo and prevents false positives in nano- diagnostics, establishing a critical new standard for high-fidelity biomarker discovery and nanomedicine.

## Introduction

The application of nanoparticles (NPs) in medicine has revolutionized the landscape of drug delivery, imaging, and diagnostics.(1–3) Upon exposure to physiological fluids, the pristine surface of a NP—its “synthetic identity”—is rapidly coated by a dynamic layer of biomolecules, dominated by proteins.(4, 5) This interface, known as the “protein corona,” confers a new “biological identity” to the NP, which ultimately dictates its interaction with cells, biodistribution, pharmacokinetics, and toxicity profiles.(6, 7) Consequently, accurate and precise characterization of the protein corona is not merely an analytical goal; it is a fundamental requirement for translating nanotechnologies from the bench to the clinic.(8, 9)

Beyond their role in therapeutic delivery, NPs are increasingly utilized as diagnostic tools. Their tunable physicochemical properties enable them to function as “scavengers” in complex biological fluids, enriching low-abundance proteins from the plasma that may serve as biomarkers for early disease detection.(10) In this diagnostic context, the protein corona effectively becomes the sample of interest. Accordingly, it is essential to identify which proteins adsorb to the NP surface. If corona profiles are confounded by non-specific carryover or co-isolated material(11, 12)

Despite the critical importance of corona characterization, commonly used isolation techniques remain a source of significant analytical variability. The most ubiquitous methods for separating corona-coated NPs from unbound plasma components or cell secretome are centrifugation and magnetic separation.(13) Although effective for bulk separation, these methods do not inherently distinguish between proteins that are truly adsorbed to the NP surface from those associated with other particulate matter present in biological fluids (e.g., cytosolic proteins from cells)(14).

Human plasma is a heterogeneous matrix that contains not only soluble proteins but also protein aggregates, lipoproteins, and extracellular vesicles (EVs), including exosomes and microvesicles.(11, 13, 15) EVs carry diverse proteins, often incorporating plasma membrane proteins via selective cargo-loading mechanisms while containing relatively few intraluminal soluble proteins.(15) Many cytosolic proteins detected in EV preparations are not encapsulated but instead externally associated, likely originating from copurifying cellular debris and aggregates.(14, 15)

EVs spanning ∼30 nm to several microns in diameter, can become trapped among engineered NPs during collection from plasma—e.g., by centrifugation or, for magnetic NPs, magnetic separation. This entrapment risks co-isolation, whereby EVs and their protein-rich cargo are unintentionally collected with the NPs. Recent evidence further suggests that protein aggregates and EVs may not only co-isolate, but also adhere to NP surfaces or compete for surface area, thereby altering the corona composition.(11, 13) Despite these concerns, rigorous quantification of this “EV effect” remains limited. As a result, it is still unclear what fraction of the identified “corona” is derived from soluble plasma proteins versus EV-associated proteins, a distinction that is essential for safety profile assessment, prediction of pharmacokinetics and biological fates of NPs and the discovery of biomarkers.

The soluble fraction of the plasma proteome represents a rich reservoir of pathophysiological information, distinct from the vesicular cargo of EVs. This “soluble” compartment is not merely a collection of passive bystanders like albumin and immunoglobulins; it contains the secretome— a vast array of proteins actively released by cells via non-vesicular mechanisms, including cytokines, growth factors, hormones, and shed receptors.(16, 17) These secreted proteins often serve as the early indicators of cellular distress, immune activation, or malignant transformation, making them highly sought-after biomarkers for conditions ranging from cardiovascular disease to cancer.(18, 19) Because of their low abundance, they are at the risk of being masked by higher abundant proteins and/or carryover material from other sources(20) Consequently, the ability of NPs to specifically enrich these low-abundance soluble markers while minimizing co-isolation of EV-associated proteins is paramount for realizing the full potential of the “protein corona” as a next-generation diagnostic tool.

In this study, we employed a rigorous comparative approach by incubating NPs in pooled human plasma versus the same plasma after depletion of an EV-enriched sedimentable fraction. EV depletion was performed by ultracentrifugation at 100,000 × g for 2 h at 4 °C. The resulting supernatant was used as EV-depleted plasma, whereas the resuspended pellet was characterized using the immunoaffinity MACSPlex multiplex bead platform targeting 37 known EV surface epitopes. MACSPlex was used only for post-isolation characterization and did not generate the EV-depleted plasma.(21, 22) We analyzed the protein corona formed on highly monodisperse polystyrene NPs of varying sizes (50 nm to 1000 nm), as well as magnetic beads consisting superparamagnetic iron oxide NPs (SPIONs- 1500 nm), which are widely used in industrial automated corona workflows. Quantitative mass spectrometry was used to compare corona composition across conditions to disentangle contributions of soluble plasma proteins versus vesicle-associated proteins. This study provides necessary insight into the origin of proteins assigned to corona, with implications for predictive safety assessment, biomarker discovery, and therapeutic target interpretation.

## Results and discussion

To systematically evaluate the impact of EVs on NPs protein corona composition, we designed a comparative study utilizing both standard pooled human plasma and plasma depleted of EVs. The overall experimental workflow is illustrated in **Figure 1**.

**Figure 1:**
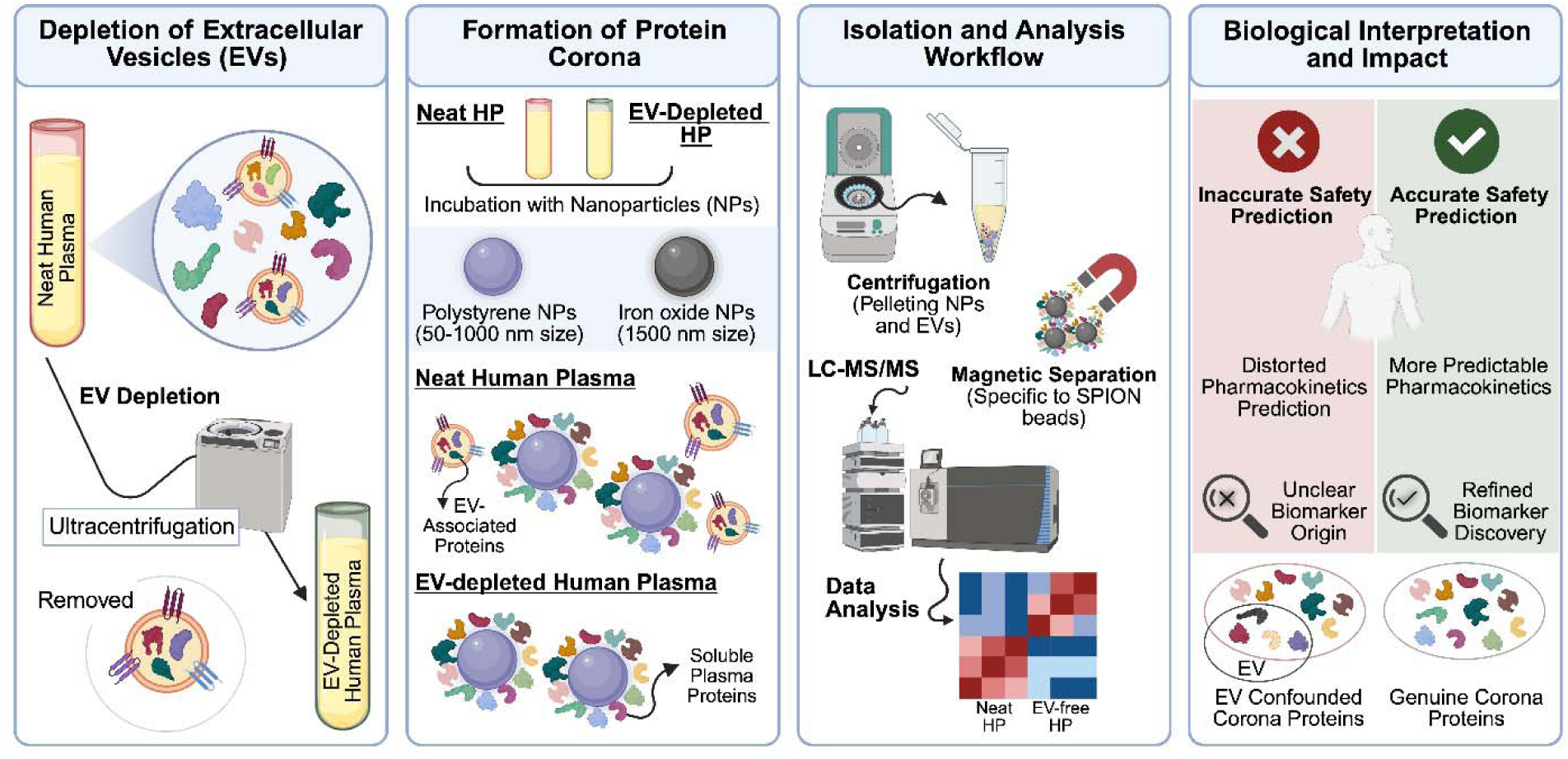
Schematic showing the overview of the workflow of this study.

A critical prerequisite for this study was to reduce EV content while retaining a plasma supernatant suitable for matched NP incubations. Pooled plasma was ultracentrifuged at 100,000 × g for 2 h at 4 °C, and the resulting supernatant constituted the EV-depleted condition. The resuspended ultracentrifugation pellet was then analyzed with the MACSPlex multiplex bead platform as a characterization assay for 37 EV surface epitopes.(21, 23) As shown in **Figure S1A**, the analysis of the captured fraction confirmed the presence of a wide variety of EV surface markers. Notably, we observed strong signals for the canonical tetraspanin exosome markers CD9, CD63, and CD81,(24) validating that the depletion process successfully targeted and removed major EV populations from the plasma. This validation ensured that the subsequent “EV-depleted plasma” condition was truly distinct from the “ regular/neat plasma” control.

Next, we incubated highly monodisperse polystyrene NPs of varying sizes (50, 100, 200, 500, 750, and 1000 nm) with either neat or EV-depleted human plasma. NP size distributions were confirmed by dynamic light scattering (DLS), as shown in **Figure S1B**. Following incubation, corona-coated NPs were recovered using a standardized centrifugation protocol, which included three rigorous washing steps to eliminate loosely bound impurities and non-specific plasma components.

As an initial qualitative assessment of the corona composition, we resolved the eluted proteins via sodium dodecyl sulfate-polyacrylamide gel electrophoresis (SDS-PAGE). The resulting protein bands provided a visual fingerprint of the corona (see **Figure S1C**) and revealed clear differences between plasma conditions. Regardless of NP size, the protein coronas formed in neat plasma displayed band patterns with distinct molecular weight distributions compared to those formed in EV-depleted plasma. Several protein bands prominent in neat plasma were either absent or significantly diminished following EV depletion. These observations suggest that a subset of proteins commonly assigned to the NP corona arises from EV carryover, either from intact vesicles retained with the particles or from EV-associated proteins and cargo that become accessible when retained vesicles are perturbed or solubilized during repeated isolation, washing, denaturation, reduction/alkylation, and enzymatic digestion rather than from proteins directly adsorbed to the NP surface.

To corroborate the biochemical results and physically locate the EVs, we employed advanced imaging techniques to visualize the NP surfaces in their native state. Cryogenic transmission electron microscopy (Cryo-TEM) was utilized to image the corona-coated polystyrene NPs suspended in vitrified ice(25) In samples incubated with neat plasma, cryo-TEM revealed a heterogeneous surface layer consistent with absorbed soluble proteins(11) alongside vesicular structures co-localized with the NPs (**Figure 2A-B**). These vesicles retained their characteristic bilayer membrane structure, confirming the presence of intact EVs within the “purified” NP pellet. Furthermore, scanning electron microscopy (SEM) provided complementary topological data, showing spherical, vesicle-like structures adhered to, or aggregated alongside, the polystyrene NPs (**Figure 2C-D**). Crucially, cryo-TEM and SEM imaging of the polystyrene NP protein corona after incubation with EV-depleted plasma revealed a complete absence of EVs (**Figure S2**). Collectively, these images provide direct visual evidence that standard centrifugation methods co-isolates EVs with PS NPs, thereby altering the protein composition recovered and subsequently assigned to the corona.

**Figure 2.**
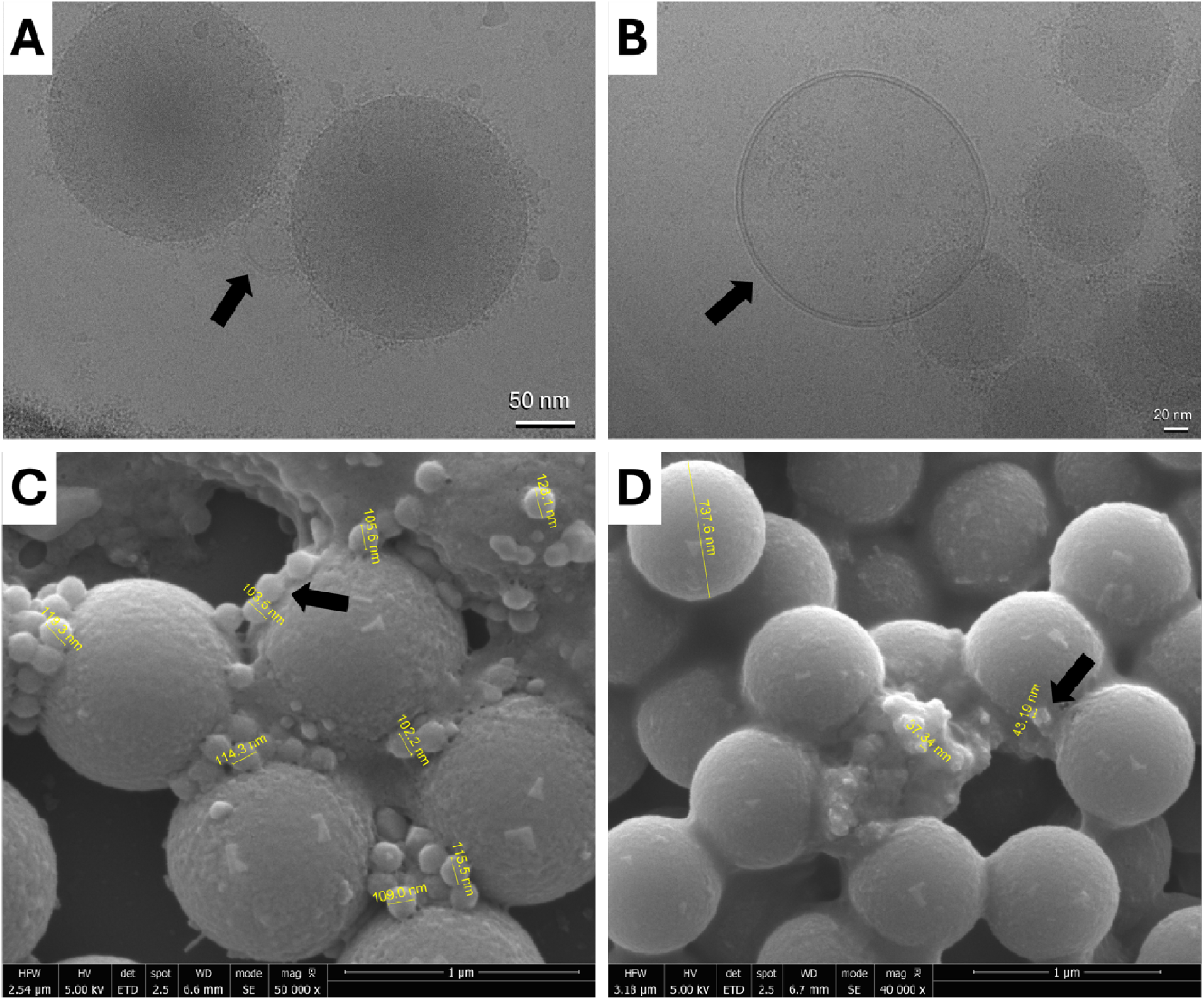
Direct visualization of EVs in protein corona. (A) and (B) Representative visualization of EV co-isolation with NPs using Cryo-TEM; (C) and (D) SEM imaging of NPs incubated in neat plasma show vesicular structures with bilayer membranes co-localized with or adhered to the NPs, providing direct visual evidence of EV presence and attachment in standard corona preparations.

To quantify the proteomic differences implied by the SDS-PAGE profiles, we analyzed recovered corona proteins by liquid chromatography-tandem mass spectrometry (LC-MS/MS). We compared the total number of identified proteins adsorbed to polystyrene NPs of varying sizes after incubation in either standard or EV-depleted plasma. As summarized in **Figure 3A**, EV removal caused a dramatic reduction in protein diversity irrespective of the NP size. In every case, the number of identified proteins dropped sharply in the EV-depleted samples compared to the standard plasma controls. For example, 50 nm NPs yielded 4,430 identified proteins in standard plasma, but only 1,139 in EV-depleted plasma, accounting for nearly a 75% reduction in protein numbers. These results indicate that a substantial fraction of proteins identified in standard corona analysis is associated with the EV-enriched fraction. Although the specific composition of the protein corona and remained size-dependent (consistent with prior reports)(5, 26, 27), the total number of identified proteins in neat plasma was largely independent of NP size, whereas the number identified in EV-depleted plasma varied with NP size.

**Figure 3.**
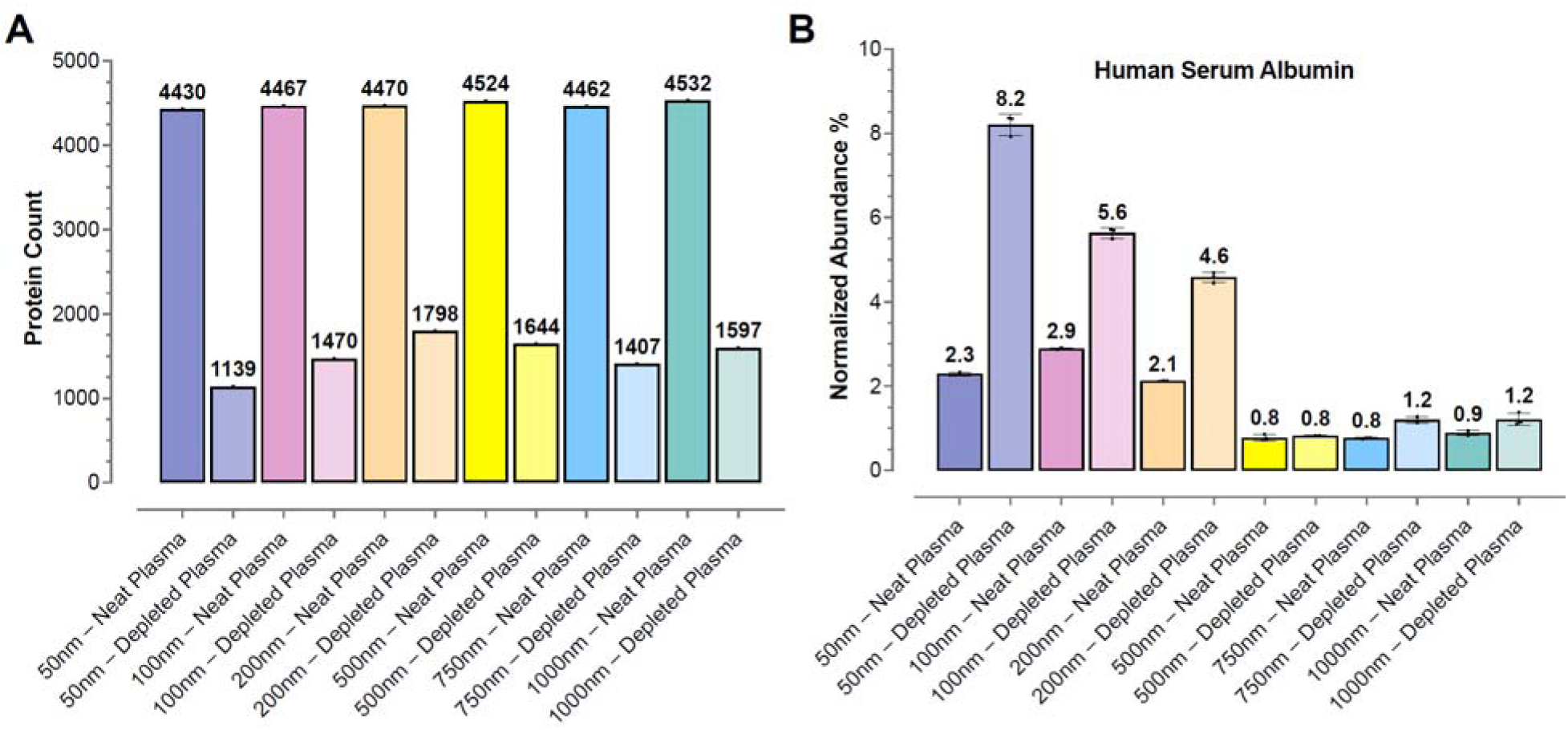
Quantitative proteomics reveals that EVs alter NP protein corona composition and reshape the representation of highly abundant plasma proteins. (A) Total count of unique proteins identified via LC-MS/MS on polystyrene NPs of difference sizes (50–1000 nm). (B) Normalized spectral abundance of albumin.

To further investigate how EVs distort the perceived composition of the corona, we specifically analyzed the behavior of human plasma albumin, the most abundant protein in plasma. Although albumin is often reported as a major component, its measured abundance varies widely across studies, suggesting sensitivity to experimental conditions and workflows. Consistent with this, we observed a clear “unmasking” effect upon EV depletion (**Figure 3B**). For smaller NPs (50–200 nm), the normalized abundance of albumin increased significantly in EV-depleted plasma compared to neat plasma. For example, on 50 nm NPs, albumin rose from 2.3% in neat plasma to 8.2% after EV-depletion. This suggests that in standard preparations, the extensive contribution from EV-associated proteins dilutes the signal of genuine corona proteins such as albumin, leading to an underestimation of their true surface coverage. In contrast, for larger NPs (500–1000 nm), albumin abundance remained relatively low (<1.5%) in both conditions, potentially indicating distinct surface curvature effects (curvature-dependent packing) or binding kinetics that minimize albumin adsorption on larger surfaces regardless of EV presence.

We next examined the rank order of albumin within the identified proteome as an orthogonal measure of enrichment depth (**Figure S3**). Rank is informative because it reflects how well the NP surface enriches specific proteins relative to the background. Consistent with EV-driven masking event, albumin ranked higher (i.e., had a lower numerical rank value, indicating higher abundance) in EV-depleted plasma as compared to neat plasma. This shift was most pronounced for larger NPs. For 500 nm NPs, albumin moved from 29th in neat plasma to 20th after EV-depletion; similarly, for 1000 nm NPs, the rank improved from 25th to 17th. These changes indicate that EV associated proteins inflate the apparent corona proteome and lower the percentage contribution of genuine corona proteins, which could potentially limit detection and quantification of soluble plasma proteins. More significantly, a population of lower abundant proteins become visible in EV-depleted samples. For example, CD27—a transmembrane glycoprotein belonging to the TNFR superfamily—was detected exclusively in EV-depleted preparations, regardless of NP size. Notably, CD27 possesses a well-characterized soluble ectodomain (sCD27) that is proteolytically shed from activated lymphocytes and circulates independently of the membrane-bound receptor, and has itself been studied as a circulating biomarker in lymphoproliferative and autoimmune disease.(28) Because peptide-level LC-MS/MS cannot distinguish the shed soluble ectodomain from the full-length protein, its selective detection here is consistent with genuine low-abundance soluble-protein unmasking rather than co-isolated cellular debris. EV-depletion therefore yields a more accurate representation of the soluble plasma proteins’ concentration at the NP surface, validating that standard isolation methods obscure the true biological identity of the NPs.

To visualize the specific compositional shifts in the “hard corona,” we generated a heatmap of the top 10 most abundant proteins identified across all NP sizes and plasma conditions (**Figure 4A**). This analysis revealed distinct protein signatures between standard and EV-depleted plasma. In neat plasma, cytoskeletal proteins such as Actin (cytoplasmic 1), Myosin light polypeptide 6, Filamin-A, and Tropomyosin alpha-4 chain consistently appeared in the top 10, particularly for smaller NPs (50–100 nm). Since these are intracellular structural proteins not typically abundant in soluble plasma,(29, 30) their presence suggests that cytosolic cargo from co-isolated EVs contributes to the recovered NP-associated protein fraction. In contrast, in the EV-depleted samples, these cytoskeletal markers(30) were largely absent from the top 10. The corona profile for EV-depleted samples shifted toward classical soluble plasma proteins, including Apolipoproteins (A-I, A-II, B-100, C-III), Clusterin, and Complement C3. The apolipoprotein signals may reflect freely exchangeable proteins, directly adsorbed proteins, and/or associated lipoprotein particles; the present workflow does not distinguish among these sources. Notably, Albumin remained a top component across most depleted samples but was often displaced by cytoskeletal carryover proteins in neat plasma. This heatmap highlights a clear fingerprint difference between the two conditions: the standard plasma-derived corona contains both NP-associated proteins and EV-associated cytosolic/structural proteins, whereas EV depletion yields a less EV-dominated profile enriched in soluble and lipoprotein-associated proteins plasma components.

**Figure 4.**
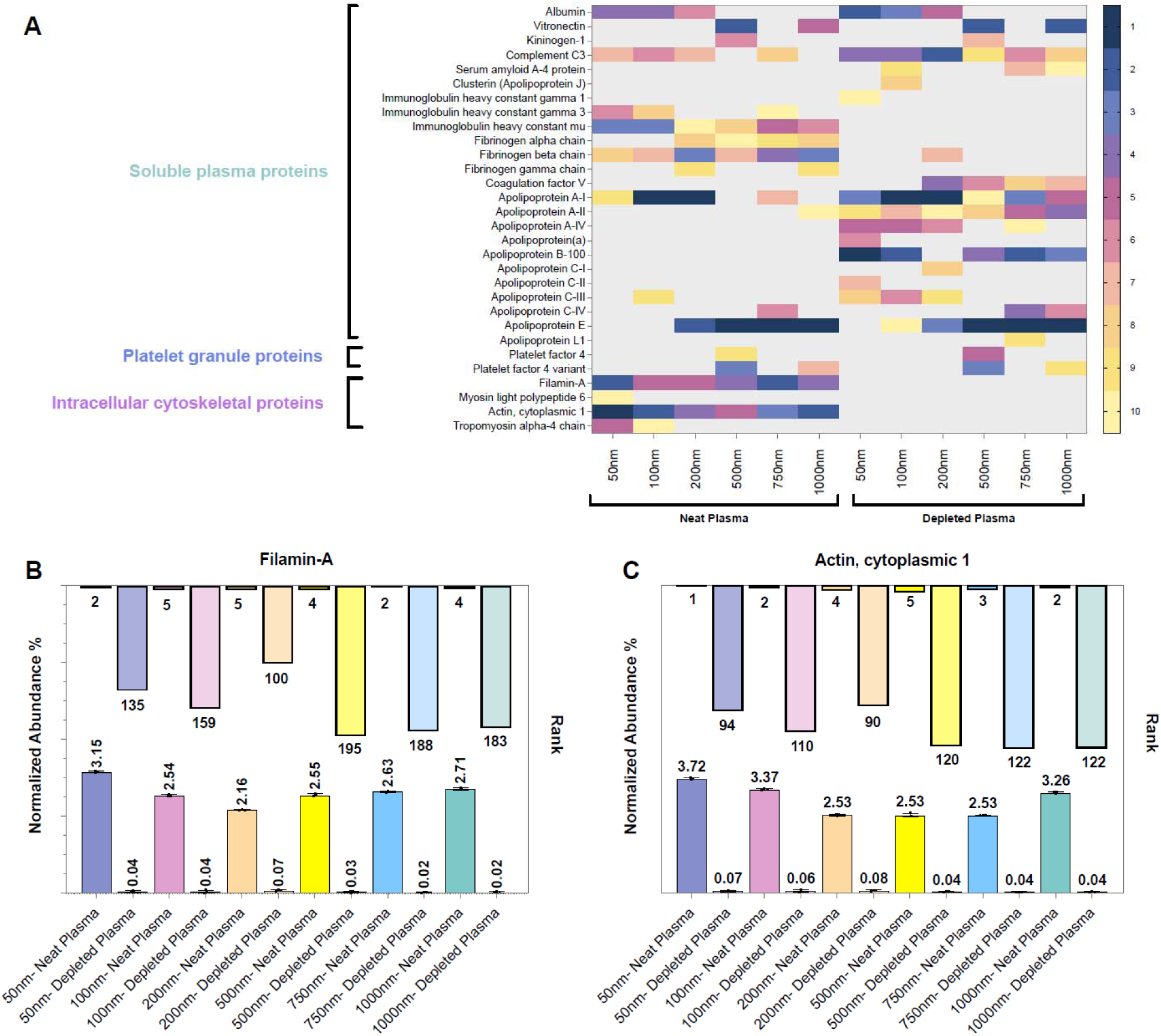
Normalized abundance and rank of intracellular cytoskeletal proteins across neat plasma and EV-depleted plasma samples in the presence of polystyrene NPs of varying sizes. Shown are (A) Heatmap of the top 10 most abundant proteins identified in each condition (Rank 1 = Dark Blue; Ran 10 = Yellow), (B) Filamin-A, (C) Actin, cytoplasmic 1. Bars along the lower x-axis represent the normalized abundance (%) of each protein under the indicated conditions, whereas bars along the upper x-axi indicate the corresponding protein rank within the NP corona.

A more detailed analysis of cytoskeletal intracellular proteins appearing among the top 10 in neat plasma samples revealed that these proteins exhibit higher ranks (i.e., lower relative abundance) in the NP corona compared with neat plasma (**Figure 4A**). This trend was observed across all NP sizes, examples of which are shown for Filamin-A and Actin (**Figure 4B-C**).

This observation prompted a more detailed examination of the hard corona composition across all experimental conditions. We first identified proteins that were consistently detected across all samples, independent of NP size and EV presence, yielding a total of 787 common proteins. To assess how the abundance of each protein varied across conditions, zero-scoring normalization was applied, and the results were visualized using a heat map. As shown in **Figure 5**, the overall abundance patterns differ markedly between neat plasma and EV-depleted plasma for the majority of these proteins.

**Figure 5.**
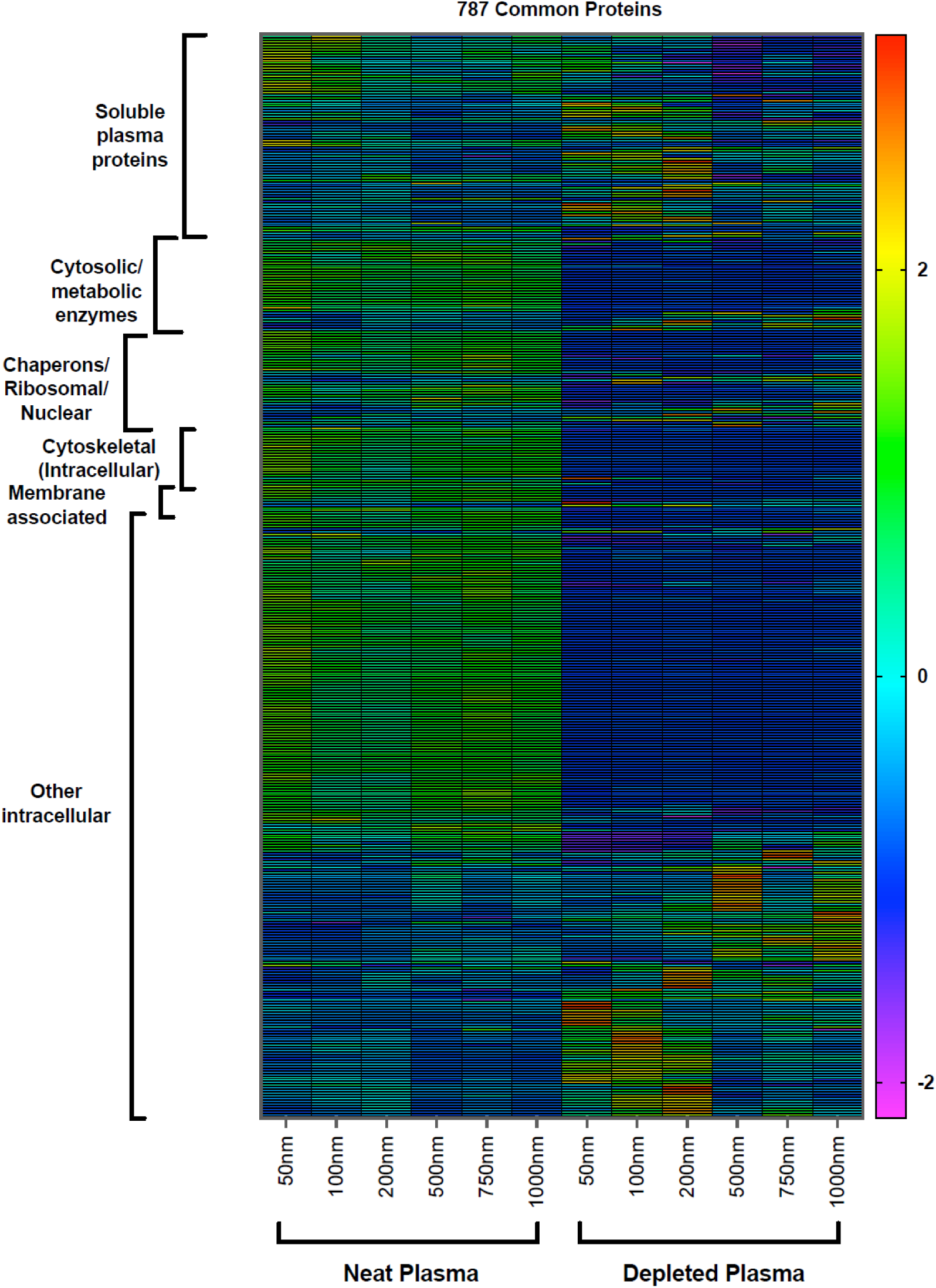
Heat map of z-scored protein abundance for proteins common to all experimental conditions. Proteins detected across all samples, irrespective of NP size or EV presence, were identified and their normalized abundances were z-scored across conditions. Rows represent individual proteins and columns represent experimental conditions. Proteins are ordered by functional category and within each category by similarity in abundance patterns. Color intensity indicates relative abundance, with positive and negative z-scores representing enrichment or depletion relative to the mean abundance of each protein across conditions.

To assess the generalizability of these observations across NP platforms and isolation strategies, we performed an analogous proteomics analysis using superparamagnetic iron oxide NP (SPION) beads, which are widely used in industrial workflows because they enable rapid particle recovery.(31, 32) We compared corona proteins isolated by both magnetic separation and standard centrifugation from neat and EV-depleted plasma. Among the many available SPION formats, we selected amine-functionalized ∼1500 nm beads because this size is representative of the magnetic bead formats commonly used in industrial and automated plasma-proteomics workflows(33–35), rather than to size-match our polystyrene NP series. Importantly, the SPION beads experiments were designed to evaluate within-platform differences between neat and EV-depleted plasma, rather than to enable direct cross-material comparisons of absolute protein counts. The total number of proteins identified on SPIONs depends strongly on particle size and surface chemistry (including coating and functional groups) and is therefore not the primary endpoint of this analysis. Although the PS NPs and SPIONs are chemically distinct, both present interfaces capable of associating with lipid-bilayer vesicles through hydrophobic, electrostatic, van der Waals, and plasma-protein/lipid bridging interactions. Magnetic bead surface chemistries are, in fact, deliberately used to capture intact EVs from plasma(32). Thus, magnetic responsiveness of the bead does not establish the molecular origin of every protein recovered with it.

Consistent with the results for polystyrene NPs, EV depletion significantly reduced the total number of identified proteins on magnetic beads (**Figure S3A**). In neat plasma, both magnetic separation and centrifugation yielded a high protein count (1,697 and 1,712 proteins, respectively). However, upon EV depletion, protein identification dropped to 904 proteins for magnetic separation and 868 proteins for centrifugation, an approximately 45-50% reduction in protein identifications. The latter data confirms that SPIONs-based workflows also suffer from substantial EV carryover, regardless of the isolation method. The similar protein counts between magnetic separation and centrifugation suggest that neither method provides a ‘cleaner’ corona profile in the presence of EVs. Albumin remained a minor component of the SPION bead corona across all conditions, with normalized abundance ranging from 1.3% to 1.7% (**Figure S3B**) with a slight increase in albumin abundance in EV-depleted conditions (e.g., 1.6% in neat magnetic vs. 1.7% in depleted magnetic). Albumin’s rank within the SPION corona proteome showed modest sensitivity to EV depletion (**Figure S3C**). Although SPIONs exhibited lower EV carryover than polystyrene NPs, this cross-platform difference cannot be assigned to a singular difference because particle size, composition, surface functionalization, and recovery physics all differ. One plausible contributor is the isolation step: polystyrene NPs were recovered at 20,000 × g for 30 min, which can sediment larger EVs and trap EVs within NP pellets, whereas magnetically recovered SPIONs are collected without exposing the bulk sample to centrifugal force. Surface-mediated EV attachment can nevertheless persist during magnetic collection, explaining why substantial EV-associated signal remained.

The heatmap of the top 10 most abundant proteins identified on SPION beads (**Figure S3D**) provided further insights. In contrast to polystyrene NPs, the predominant proteins on SPIONs were consistently dominated by highly abundant plasma proteins involved in coagulation (e.g., Fibrinogen alpha, beta, and gamma chains, Prothrombin), adhesion (vitronectin), complement activation (C4b-binding protein alpha chain, Complement C4-A), and general immunity (Immunoglobulin heavy constant gamma and mu chains, Fibronectin). These results highlight that EV carryover is a general issue that extends to industrially relevant NP platforms, and that standard isolation methods (magnetic or centrifugation) are equally susceptible to co-isolation artifacts.

Given the increasing use of NPs for biomarker discovery, we evaluated the presence of 227 FDA-approved and diagnostic plasma biomarkers within the protein coronas formed under all experimental conditions for polystyrene NPs (**Table S1**). From this list, 94 biomarkers were consistently detected across all conditions and 39 of them were detected at least in one condition as depicted in a binary heatmap in **Figure 6A**. Notably, 13 biomarkers were detected in neat plasma coronas but were absent under EV-depleted conditions suggesting EV-dependent enrichment. Additionally, detection of some biomarkers seems to be size dependent. For example, Tartrate-resistant acid phosphatase type 5 is detected at NP sizes greater than 500 nm.

**Figure 6.**
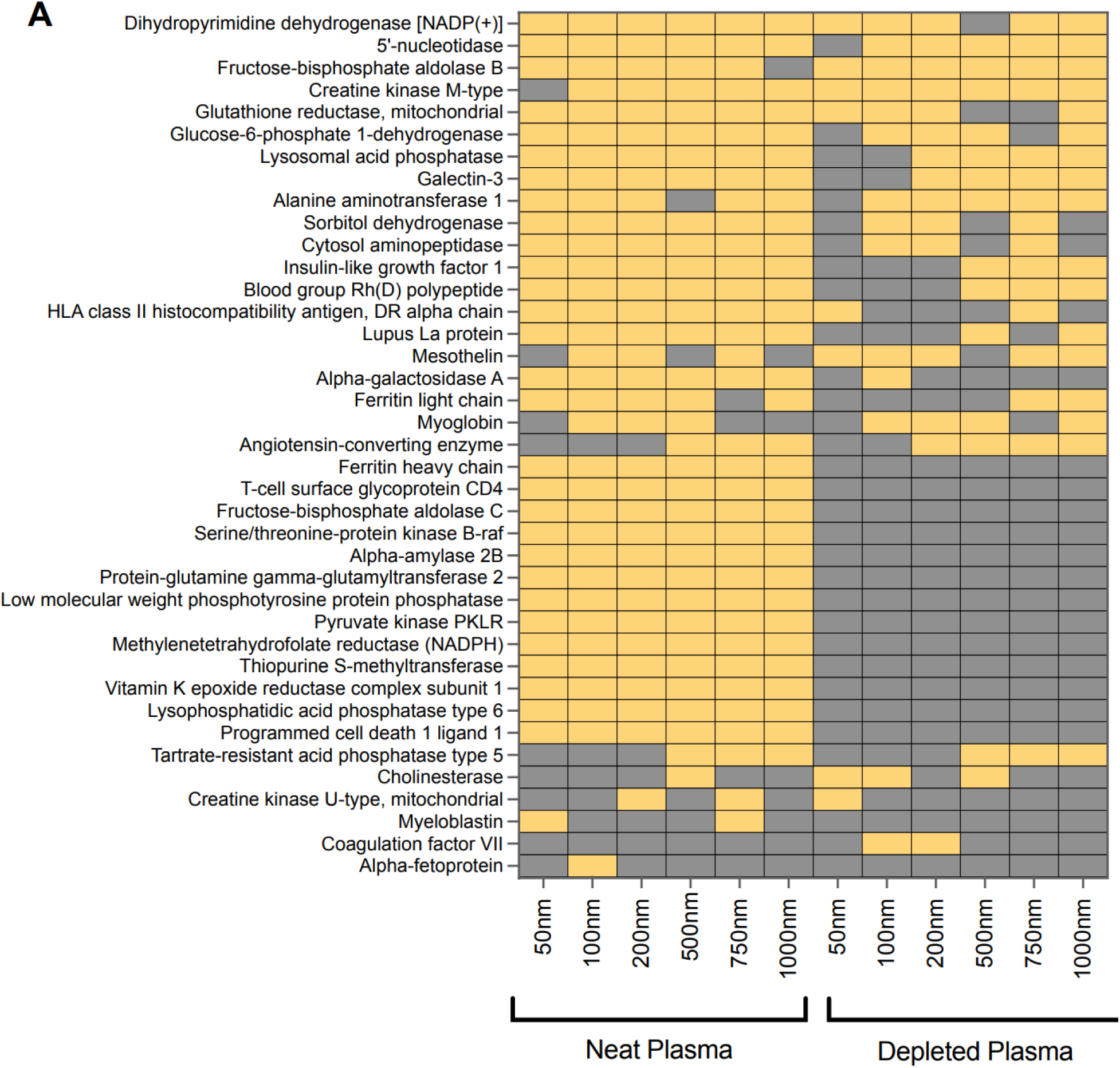
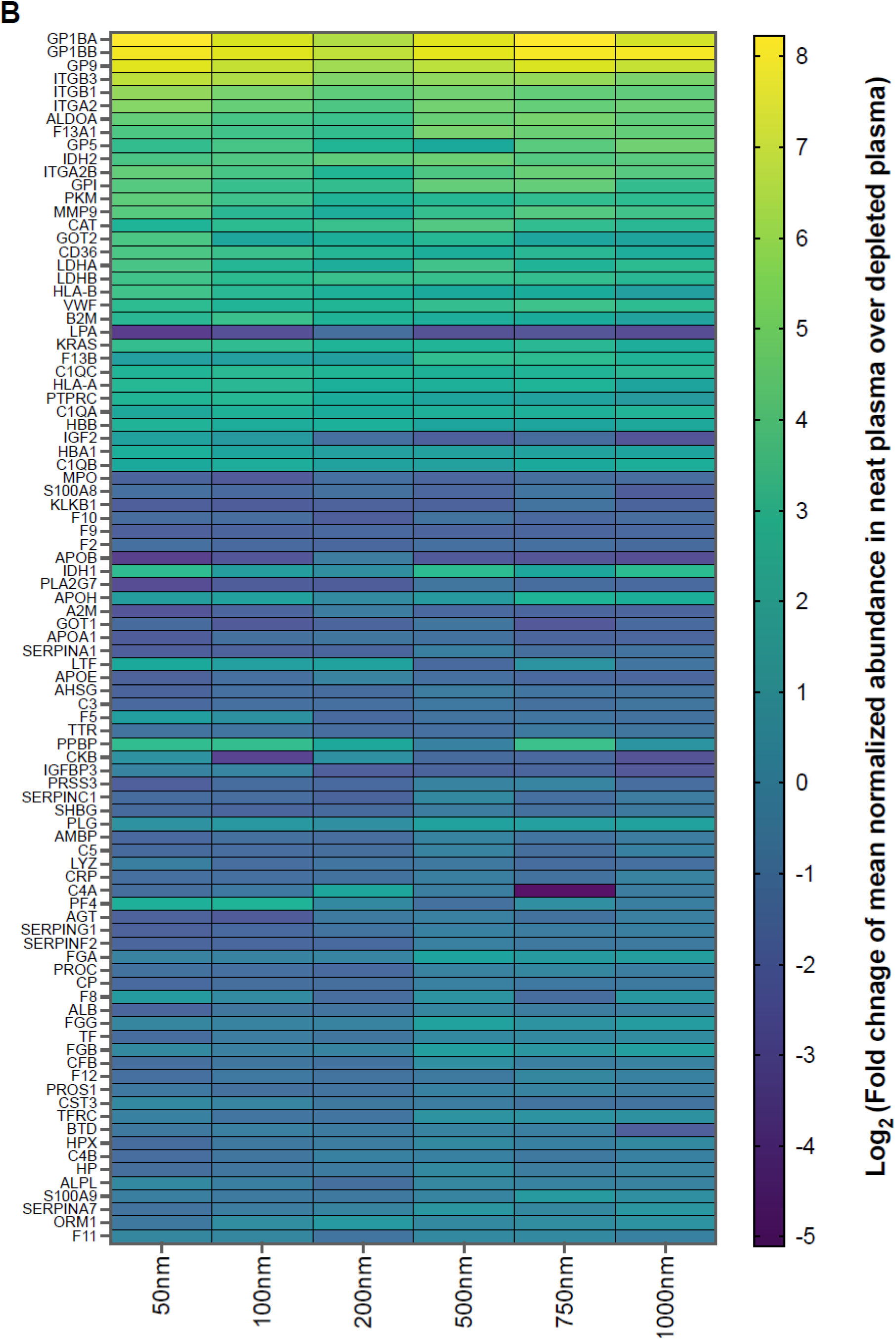
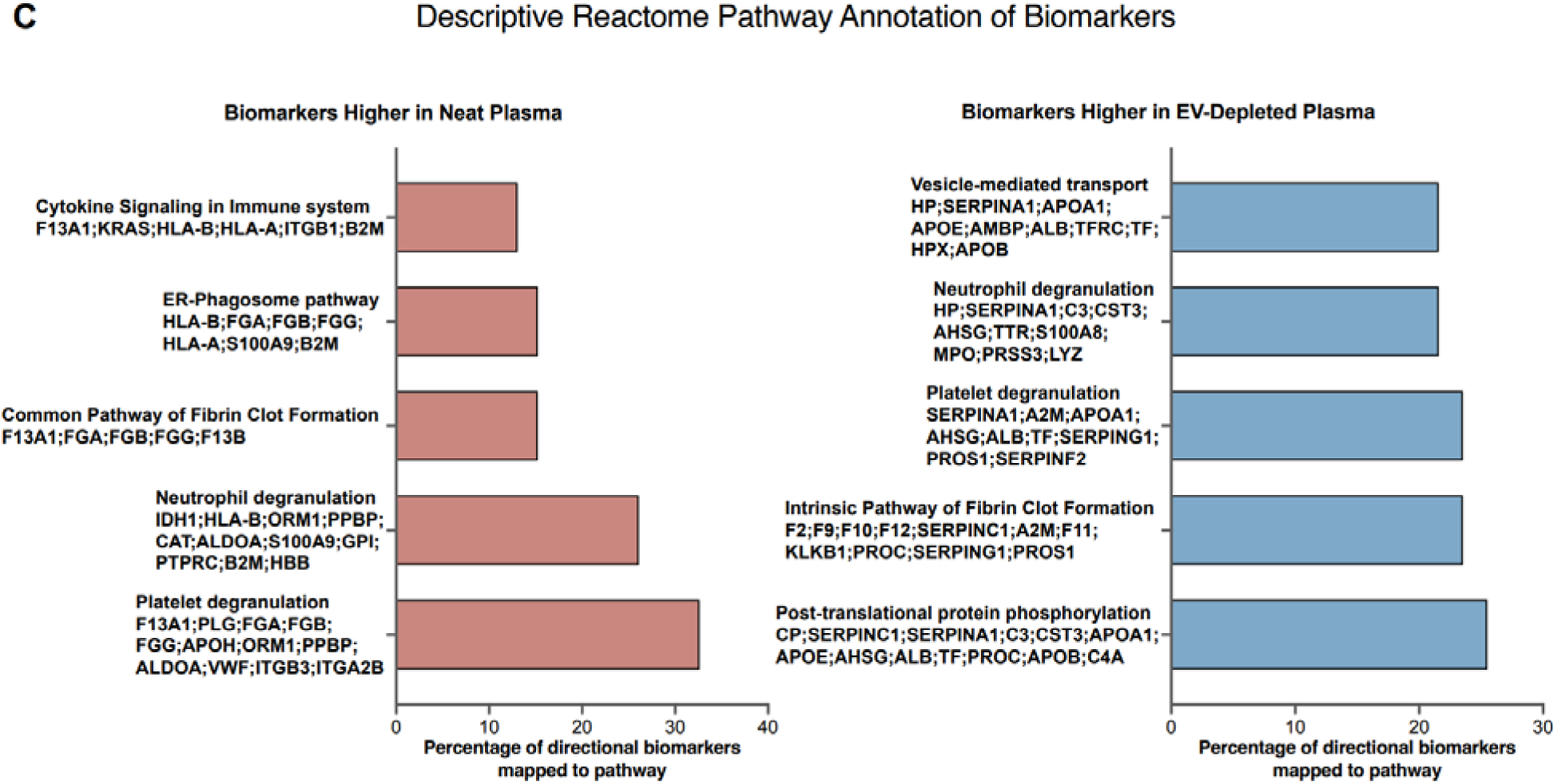
EVs’ influence on detection of FDA-approved and clinical biomarkers in protein coronas of PSNPs of different sizes. (A) Heatmap illustrating the detection (yellow) or absence (gray) of 39 FDA-approved and diagnostic plasma biomarkers in the protein corona of polystyrene NPs across various sizes (50-1000 nm). The horizontal axis represents different NP sizes and plasma conditions (neat plasma vs. depleted plasma). (B) Heatmap showing the Log_2_ of the fold change of the normalized abundance in neat plasma over EV-depleted plasma for FDA-approved and clinical plasma biomarkers commonly detected across nanoparticle protein coronas. For each nanoparticle size, replicate measurements were averaged prior to calculation. Positive values indicate higher enrichment in neat plasma, whereas negative values indicate higher enrichment in EV-depleted plasma. (C) Descriptive Reactome pathway annotation of biomarkers affected by EV depletion. Biomarkers showing at least a twofold difference in normalized nanoparticle-corona abundance between neat and EV-depleted plasma in one or more nanoparticle sizes were separated according to the direction of change. Proteins showing meaningful differences in opposite directions across nanoparticle sizes were excluded from the directional analysis. Bars represent the percentage of biomarkers within each directional list that mapped to the indicated Reactome pathway, and the contributing gene symbols are shown beside each pathway.

To further quantify differences in biomarker enrichment between neat and EV-depleted plasma for biomarkers commonly detected across conditions (94 proteins), the mean normalized abundance of each biomarker in neat plasma was compared with that in EV-depleted plasma using log2 fold change. The resulting log2 fold-change values for all commonly detected biomarkers are displayed as a heat map (**Figure 6B**). Positive values indicate higher enrichment in neat plasma coronas, whereas negative values indicate higher enrichment in EV- depleted plasma coronas. by differences in absolute protein abundance.

Analysis of the magnetic bead samples identified 88 FDA-approved and diagnostic biomarkers common to all conditions, with 17 detected only in at least one subset (**Figure S4A**). Out of these proteins, 10 present in neat plasma were absent following EV depletion.

The log2 fold-change analysis was also performed for SPION coronas separated using both magnetic separation and centrifugation methods (**Figure S4B**). Overall, biomarker enrichment patterns were largely consistent between SPION recovery methods, indicating that magnetic separation and centrifugation did not substantially alter the relative enrichment trends. However, method-dependent variation was observed for a small subset of biomarkers, most notably Integrin beta-3 and Integrin alpha-IIb, which showed log2 fold-change values of approximately 6.

These findings carry important implications for NP-based diagnostics. Studies employing NP enrichment for biomarker discovery without adequately addressing the potential for EV carryover may risk reporting false positives or misattributing the biological origin and significance of identified biomarkers. This would be of value for researchers that want or need to know the origins of the biomarker, namely plasma or EV-associated, as opposed to seeing all biomarkers, irrespective of the source. The apparent presence or absence of these crucial biomarkers in the corona is dramatically altered by EVs, unequivocally demonstrating that a cleaner, EV-depleted corona preparation is essential for accurate and reliable biomarker identification and validation.

A striking, concrete illustration of this diagnostic risk has since emerged in Alzheimer’s disease plasma proteomics using this same corona platform. Two plasma proteomic studies of Alzheimer’s disease reported opposite directions of change for CALCOCO1: a solution-phase immunoassay platform reported it as decreased in disease(36), whereas our nanoparticle corona platform reported it as increased(37). The most parsimonious explanation is that CALCOCO1 is substantially an EV-cargo protein, such that its apparent abundance, and even the direction of its disease-associated change, depends on whether a given analytical platform captures the vesicular compartment or is instead restricted to the free, soluble fraction. Analyzing our proteomics results here, we noticed that CALCOCO1, a cytoplasmic reticulophagy receptor lacking both a transmembrane domain and a conventional secretory signal peptide, was consistently detected in the corona formed on neat human plasma across all polystyrene NP sizes tested, but was undetectable or markedly reduced in the corona formed on EV-depleted plasma from the same samples (**Figure 7**). This example demonstrates precisely the diagnostic and therapeutic risk our corona data predict: for any candidate biomarker whose plasma pool is substantially EV-derived, failing to resolve its soluble-versus- vesicular origin can not only inflate false-positive rates but invert the reported direction of a disease association, undermining diagnostic interpretation and any therapeutic strategy premised on that biomarker tracking a specific, defined fraction of the total protein pool. The compartmental origin of an Alzheimer’s disease–associated protein is also a pharmacological variable, not merely an analytical question. A soluble plasma protein is generally accessible to circulating antibodies or pharmaceuticals, whereas a protein enclosed within an EV particle is shielded by the lipid bilayer and may require EV-directed delivery to be engaged.

**Figure 7.**
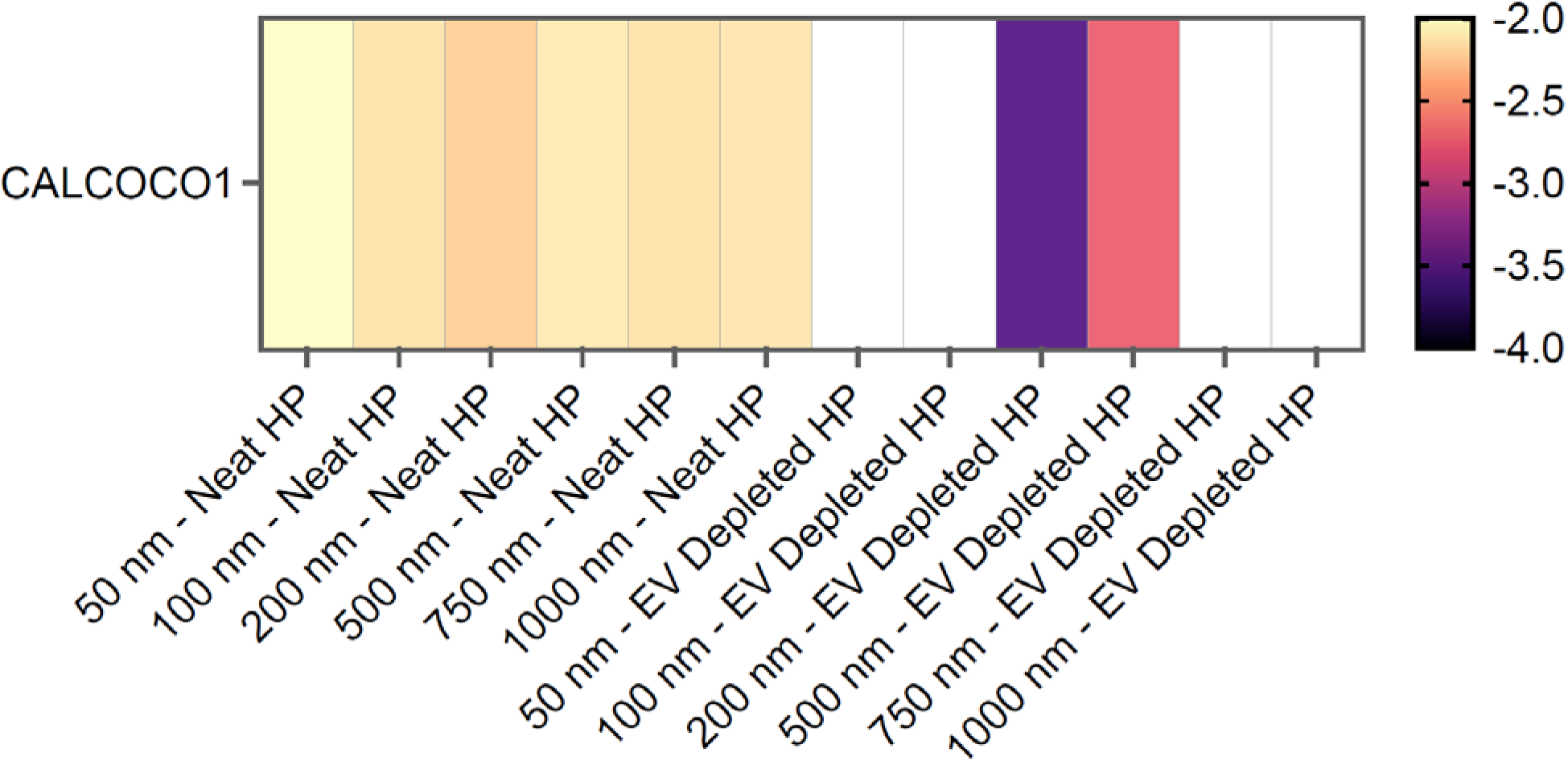
CALCOCO1 detection in the NP protein corona is lost upon EV depletion. Heatmap of normalized CALCOCO1 abundance (color scale; more negative values indicate lower abundance) across all polystyrene NP sizes (50–1000 nm) in neat plasma (left) and EV-depleted plasma (right). CALCOCO1 was detected at consistent abundance across all NP sizes in neat plasma, but was undetectable (blank cells) in EV-depleted plasma for most NP sizes, and markedly reduced in abundance at the two sizes (200 and 500 nm) where it remained detectable.

It is important to acknowledge that EVs themselves harbor a rich repertoire of proteomic information and are increasingly recognized as potent circulating biomarkers.(32, 38, 39) Consequently, in broad-spectrum plasma proteomics where the goal is to catalogue the entirety of the circulating proteome, the presence of EVs is not a “contamination” but a valuable signal. However, in the specific context of NP-enabled proteomics, the inadvertent and indiscriminate co-isolation of EVs presents a fundamental analytical challenge. Crucially, not all EVs present in a plasma sample reflect the patient’s physiological state; a significant portion can be artifacts of the blood collection process itself.(40) Standard phlebotomy and sample handling can induce platelet activation and cell lysis, releasing a flood of non-physiological EVs and granular proteins into the sample.(41, 42) These ex vivo artifacts, which do not exist in circulation, can disproportionately dominate the NP corona, effectively drowning out genuine low-abundance soluble biomarkers. Furthermore, this co-isolation obscures the biological origin of the signal; it becomes impossible to discern whether a detected biomarker was scavenged from the soluble phase by the nanoparticle’s surface chemistry or merely brought down as a passenger on a co-sedimenting, potentially artifactual, vesicle. Therefore, to achieve high-precision diagnostics, it is not that EVs should be ignored, but rather that the soluble corona must be deconvoluted from both endogenous vesicular cargo and pre-analytical contaminants.(40, 43)

These findings have broad implications for both nanomedicine and NP-enabled diagnostics. Firstly, in nanomedicine, the “biological identity” of a NP, which governs its interactions with cells, biodistribution, and ultimately its safety and efficacy, has likely been misinterpreted.(9, 44–46) If a significant portion of the “corona” is, in fact, EV cargo, then previous studies on cellular uptake mechanisms, immune responses, and pharmacokinetic profiles may need re-evaluation.(47–49) The true corona (i.e., proteins genuinely adsorbed to the NP surface) could be far simpler and functionally distinct from what was inferred from samples affected by EV carryover(40) If NPs co-enrich EV-associated proteins alongside genuine soluble biomarkers, distinguishing meaningful signals becomes incredibly challenging. By providing an approach that minimizes EV carryover, our work provides a strategy to obtain a cleaner, more accurate corona profile, paving the way for more precise biomarker identification.

## Conclusion

This study highlights a critical and often underappreciated challenge in NP research: the pervasive influence of EVs on protein corona analysis. Beyond the inherent presence of EVs in biological fluids, our findings underscore the critical importance of standardized pre-analytical sample handling; phlebotomy procedures themselves can inadvertently induce platelet activation and the subsequent release of additional, non-physiological EVs, further compromising the integrity of protein corona analysis.(40, 50, 51) Collectively, our data show that conventional workflows for isolating corona-coated NPs routinely co-isolate EVs, thereby creating a complex, artifact-ridden “biological identity” that masks the true interactions between NPs and soluble plasma proteins. In contrast, ultracentrifugal depletion of an EV-enriched sedimentable fraction reveals a less complex recovered proteome with greater relative representation of soluble plasma proteins. This deeper understanding is not merely academic refinement; it is a transformative insight for the development of safe and effective nanomedicines and the high-fidelity discovery of disease biomarkers.

For nanomedicine, these results suggest that long-standing interpretations of NP biodistribution, cellular uptake, and immunogenicity may rest on an incomplete picture of the recovered material. The accurate determination of a NP’s biological identity is significantly impacted by the composition of its protein corona. In plasma, NPs encounter soluble proteins and EVs, and standard corona recovery methods can co-purify both. A critical issue arises during mass spectrometry analysis, where the protein content of EVs and protein aggregates is erroneously included in the characterization of the protein corona. Since the biological response to NPs is highly dependent on the specific conformation, composition, and surface presentation of these proteins, the resulting mass spectrometry data, which represents an inseparable mixture of EV and soluble protein coatings, leads to an inaccurate portrayal of the NP’s true biological identity. The “corona” that dictates these outcomes is far more nuanced, demanding re-evaluation of how we predict and control NP-biological interactions. It compels us to move beyond simply identifying proteins and to ensure we are identifying the *right* proteins, those genuinely adsorbed to the NP surface.

For biomarker discovery, our findings serve as a powerful cautionary message and a clear path forward. NPs, touted as “scavengers” for low-abundance biomarkers, also inadvertently become “collectors” of EV-derived proteomic noise. This noise is compounded by artifactual EVs introduced during plasma sample collection.(52) Without explicitly accounting for EV carryover, NP-enrichment datasets risk elevated false-positive rates and ambiguous biological interpretation. In contrast, distinguishing the true corona constituents from this vesicular masquerade yields a high-fidelity readout that can unlock higher-resolution biomarker signals and accelerate the translation of NP-based diagnostics into clinical reality. The same distinction is essential for therapeutics, where soluble, EV-surface, and intravesicular pools of a candidate target can differ substantially in drug accessibility even when they contribute to the same total plasma-protein measurement.

In essence, our work compels a paradigm shift in protein corona research: from merely observing the apparent “biological identity” to actively defining its *true* composition. This crucial step is essential to harness the full potential of NPs, ushering in an era of more predictable nanotherapeutics and more precise nano-diagnostics that truly reflect the intricacies of biological systems.

## Data availability

The proteomic data have been deposited to the ProteomeXchange Consortium (https://www.proteomexchange.org/) via the MassIVE(53) partner repository (https://massive.ucsd.edu/) with MassIVE data set identifier MSV000100960 and ProteomeXchange identifier PXD074849. Excell files of the mass spectrometry outcomes are available in the Datasets 1 (for polystyrene NPs) and 2 (for magnetic beads).

## Competing Interest Statement

M.M. is a co-founder and director of the Academic Parity Movement (www.paritymovement.org). He is a co-founder of XProteome, Targets’ Tip and AlbuDerm, and he receives royalties/honoraria for his published books, plenary lectures and licensed patents. A.A.S. and B.B. are co-founders of XProteome.

## Supporting Information

The experimental methods for the preparation of the protein corona, as well as the mass spectrometry analysis, are detailed in the SI.

## Notes

### Summary of Updates

Added a new dataset (Figure 7) showing an example of detecting an EV-packed protein (CALCOCO1) using our proposed platform. Also improving method section for the EV removal of depleted plasma.

## References

1. B. Y. Kim, J. T. Rutka, W. C. Chan, Nanomedicine. N Engl J Med 363, 2434–2443 (2010).

2. S. M. Moghimi, A. C. Hunter, J. C. Murray, Nanomedicine: current status and future prospects. Faseb j 19, 311–330 (2005).

3. B. Pelaz et al., Diverse Applications of Nanomedicine. ACS Nano 11, 2313–2381 (2017).

4. T. Cedervall et al., Understanding the nanoparticle-protein corona using methods to quantify exchange rates and affinities of proteins for nanoparticles. Proc Natl Acad Sci U S A 104, 2050–2055 (2007).

5. M. P. Monopoli, C. Åberg, A. Salvati, K. A. Dawson, Biomolecular coronas provide the biological identity of nanosized materials. Nature Nanotechnology 7, 779–786 (2012).

6. K. A. Dawson, Y. Yan, Current understanding of biological identity at the nanoscale and future prospects. Nature Nanotechnology 16, 229–242 (2021).

7. M. Hadjidemetriou et al., In vivo biomolecule corona around blood-circulating, clinically used and antibody-targeted lipid bilayer nanoscale vesicles. ACS nano 9, 8142–8156 (2015).

8. M. Faria et al., Minimum information reporting in bio–nano experimental literature. Nature Nanotechnology 13, 777–785 (2018).

9. P. C. Ke, S. Lin, W. J. Parak, T. P. Davis, F. Caruso, A Decade of the Protein Corona. ACS Nano 11, 11773–11776 (2017).

10. M. Mahmoudi, M. P. Landry, A. Moore, R. Coreas, The protein corona from nanomedicine to environmental science. Nature Reviews Materials 8, 422–438 (2023).

11. S. Sheibani et al., Nanoscale characterization of the biomolecular corona by cryo-electron microscopy, cryo-electron tomography, and image simulation. Nature Communications 12, 573 (2021).

12. K. Roger et al., Mining the plasma proteome: Evaluation of enrichment methods for depth and reproducibility. Journal of Proteomics 321, 105519 (2025).

13. M. J. Hajipour et al., An Overview of Nanoparticle Protein Corona Literature. Small 19, 2301838 (2023).

14. M. Unnikrishnan, Y. Wang, M. Gruebele, C. J. Murphy, Nanoparticle-assisted tubulin assembly is environment dependent. Proceedings of the National Academy of Sciences 121, e2403034121 (2024).

15. J. Morales-Sanfrutos et al., Defining the reference proteomes for small extracellular vesicles and non-vesicular components. Nature Cell Biology 28, 622–639 (2026).

16. M. Uhlén et al., The human secretome. Science signaling 12, eaaz0274 (2019).

17. C. Gabay, I. Kushner, Acute-phase proteins and other systemic responses to inflammation. New England journal of medicine 340, 448–454 (1999).

18. V. Kulasingam, E. P. Diamandis, Strategies for discovering novel cancer biomarkers through utilization of emerging technologies. Nature clinical practice Oncology 5, 588–599 (2008).

19. N. L. Anderson, N. G. Anderson, The human plasma proteome: history, character, and diagnostic prospects. Molecular & cellular proteomics 1, 845–867 (2002).

20. P. E. Geyer et al., Plasma proteome profiling to assess human health and disease. Cell systems 2, 185–195 (2016).

21. O. P. B. Wiklander et al., Systematic Methodological Evaluation of a Multiplex Bead-Based Flow Cytometry Assay for Detection of Extracellular Vesicle Surface Signatures. Front Immunol 9, 1326 (2018).

22. A. Brahmer et al., Platelets, endothelial cells and leukocytes contribute to the exercise-triggered release of extracellular vesicles into the circulation. Journal of extracellular vesicles 8, 1615820 (2019).

23. S. Grumelot et al., Lipid nanoparticle protein coronas form via lipoprotein fusion rather than shell-like adsorption. bioRxiv 10.64898/2025.12.21.695162, 2025.2012.2021.695162 (2025).

24. M. Mathieu et al., Specificities of exosome versus small ectosome secretion revealed by live intracellular tracking of CD63 and CD9. Nature communications 12, 4389 (2021).

25. X.-C. Bai, G. McMullan, S. H. Scheres, How cryo-EM is revolutionizing structural biology. Trends in biochemical sciences 40, 49–57 (2015).

26. M. Lundqvist et al., Nanoparticle size and surface properties determine the protein corona with possible implications for biological impacts. Proceedings of the National Academy of Sciences 105, 14265–14270 (2008).

27. M. P. Monopoli et al., Physical−Chemical Aspects of Protein Corona: Relevance to in Vitro and in Vivo Biological Impacts of Nanoparticles. Journal of the American Chemical Society 133, 2525–2534 (2011).

28. W. A. Loenen et al., The CD27 membrane receptor, a lymphocyte-specific member of the nerve growth factor receptor family, gives rise to a soluble form by protein processing that does not involve receptor endocytosis. European journal of immunology 22, 447–455 (1992).

29. Y.-J. Liu, C. Wang, A review of the regulatory mechanisms of extracellular vesicles-mediated intercellular communication. Cell Communication and Signaling 21, 77 (2023).

30. D. S. Choi, D. K. Kim, Y. K. Kim, Y. S. Gho, Proteomics of extracellular vesicles: exosomes and ectosomes. Mass spectrometry reviews 34, 474–490 (2015).

31. J. E. Blume et al., Rapid, deep and precise profiling of the plasma proteome with multi-nanoparticle protein corona. Nature Communications 11, 3662 (2020).

32. C. C. Wu et al., Enrichment of extracellular vesicles using Mag-Net for the analysis of the plasma proteome. Nature Communications 16, 5447 (2025).

33. C. S. Hughes et al., Single-pot, solid-phase-enhanced sample preparation for proteomics experiments. Nature Protocols 14, 68–85 (2019).

34. T. Müller et al., Automated sample preparation with SP3 for low-input clinical proteomics. Mol Syst Biol 16, e9111 (2020).

35. S. A. Sadeghi et al., Mass spectrometry-based top-down proteomics for proteoform profiling of protein coronas. Nature Protocols 21, 1092–1125 (2026).

36. G. Bellomo et al., Plasma proteome profiling identified biomarkers for the differential diagnosis and molecular staging of neurodegenerative dementias. Nature Aging 10.1038/s43587-026-01162-7 (2026).

37. N. Mahmoudi et al., Deep nanoparticle protein corona plasma proteomics resolves a stage-specific peripheral signature of Alzheimer’s disease. bioRxiv 10.64898/2026.07.25.740710, 2026.2007.2025.740710 (2026).

38. A. Hoshino et al., Tumour exosome integrins determine organotropic metastasis. Nature 527, 329–335 (2015).

39. A. Dickhout, R. R. Koenen, Extracellular vesicles as biomarkers in cardiovascular disease; chances and risks. Frontiers in cardiovascular medicine 5, 113 (2018).

40. H. Tang, J. Wang, M. Mahmoudi, Improving accuracy and reproducibility of mass spectrometry characterization of protein coronas on nanoparticles. Nature Protocols 20, 3057–3063 (2025).

41. R. Lacroix et al., Standardization of platelet-derived microparticle enumeration by flow cytometry with calibrated beads: results of the International Society on Thrombosis and Haemostasis SSC Collaborative workshop. Journal of Thrombosis and Haemostasis 8, 2571–2574 (2010).

42. J. A. Welsh et al., Minimal information for studies of extracellular vesicles (MISEV2023): From basic to advanced approaches. J Extracell Vesicles 13, e12404 (2024).

43. M. L. Merchant, I. M. Rood, J. K. Deegens, J. B. Klein, Isolation and characterization of urinary extracellular vesicles: implications for biomarker discovery. Nature Reviews Nephrology 13, 731–749 (2017).

44. M. Faria et al., Minimum information reporting in bio-nano experimental literature. Nat Nanotechnol 13, 777–785 (2018).

45. M. Faria, S. Spoljaric, F. Caruso, Reanalysis: the forgotten sibling of reproducibility and replicability. Nature Reviews Methods Primers 2, 14 (2022).

46. S. Li, C. Cortez-Jugo, Y. Ju, F. Caruso, Approaching Two Decades: Biomolecular Coronas and Bio–Nano Interactions. ACS Nano 18, 33257–33263 (2024).

47. J. B. Simonsen, What are we looking at? Extracellular vesicles, lipoproteins, or both? Circulation research 121, 920–922 (2017).

48. J. B. Simonsen, Technical challenges of studying the impact of plasma components on the efficacy of lipid nanoparticles for vaccine and therapeutic applications. Nature Communications 15, 3852 (2024).

49. J. B. Simonsen, Lipid nanoparticle-based strategies for extrahepatic delivery of nucleic acid therapies - challenges and opportunities. J Control Release 370, 763–772 (2024).

50. F. Mullier, N. Bailly, C. Chatelain, B. Chatelain, J.-M. Dogné, Pre-analytical issues in the measurement of circulating microparticles: current recommendations and pending questions. Journal of Thrombosis and Haemostasis 11, 693–696 (2013).

51. R. Lacroix et al., Impact of pre-analytical parameters on the measurement of circulating microparticles: towards standardization of protocol. Journal of Thrombosis and Haemostasis 10, 437–446 (2012).

52. J. A. Welsh et al., Minimal information for studies of extracellular vesicles (MISEV2023): From basic to advanced approaches. Journal of Extracellular Vesicles 13, e12404 (2024).

53. M. Choi et al., MassIVE. quant: a community resource of quantitative mass spectrometry–based proteomics datasets. Nature methods 17, 981–984 (2020).

