## Supplementary material for "Extracellular Vesicle Carryover Distorts Nanoparticle Protein Corona Profiles in Human Plasma": SI

**Materials and Methods**

**Human plasma and EV depletion.** Pooled healthy human plasma was procured from BioIVT Company ([www.bioivt.com](http://www.bioivt.com/)) and stored according to manufacturer's instructions. The fasting status of the plasma donors was not specified by the vendor. Extracellular vesicles (EVs) were removed from a portion of the pooled plasma by ultracentrifugation at 100,000 × g for 2 hours at 4°C; the resulting supernatant constituted the EV-depleted plasma used for subsequent NP incubations, while the pelleted EVs were resuspended in phosphate-buffered saline (PBS) for downstream characterization.

**MACSPlex characterization of the ultracentrifugation pellet.** To confirm that this pelleted fraction consisted of genuine EVs, and to verify that the antibody-coated capture beads do not simply adsorb soluble protein nonspecifically, the resuspended EV pellet and, separately, an aliquot of neat plasma were each incubated with the MACSPlex Exosome Capture Beads Kit (Miltenyi Biotec, Cat No. 130-108-813) according to the manufacturer's protocol. This kit enables characterization of EVs via antibody capture of 37 distinct EV surface epitopes. EVs captured by the MACSPlex beads were detected using fluorescently labeled antibodies against pan-exosome markers such as CD9, CD63, and CD81. Flow cytometry (e.g., using a Miltenyi Biotec MACSQuant Analyzer) was employed to acquire semiquantitative data on the expression of the 37 EV surface epitopes. Calibration of the flow cytometer involved adjusting background settings to unlabeled beads and establishing gating strategies to identify individual bead populations for each analyte. Batch analysis quantified median fluorescence intensities for each bead population, and EV surface marker expression was calculated for each sample. As a control, and to show that the beads do not adsorb soluble proteins, 20 µL of plasma sample was incubated with the MACSPlex beads according to the manufacturer's protocol, allowing the capture of diverse EV subpopulations (including exosomes) through specific antigen-antibody interactions. The EV-depleted plasma was then collected from the supernatant after removal of the bead-captured EVs via centrifugation, ensuring that the plasma used for subsequent NP incubations was significantly depleted of endogenous vesicles while preserving its soluble proteome. The effectiveness of EV depletion was confirmed by analyzing the captured fraction.

**Nanoparticles**. Highly monodisperse carboxyl-terminated polystyrene WNPs (sizes: 50, 100, 200, 500, 750, and 1000 nm; sourced from PolySciences, Inc.) were used for the primary protein corona studies. Magnetic beads consist of superparamagnetic iron oxide NPs (SPIONs, BioMag®, average size of 1.5 µm) were also utilized to assess the generalizability of the findings and their applicability to industrially relevant magnetic particles. NPs were stored as aqueous suspensions and handled according to manufacturer’s guidelines.

**Protein corona formation and isolation**. We use our optimized protein corona formation and characterization protocols(1) prior to mixing human plasma and nanoparticles (e.g., removing protein aggregates and particulate matter(2)). For protein corona formation, NPs were mixed with either regular/neat or EV-depleted human plasma at a final plasma concentration of 55% and NPs’ concentration of 200 ng/ml. The NP-plasma mixtures were incubated at 37°C for 1 hour with constant orbital agitation to ensure homogeneous protein adsorption. Following incubation, corona-coated NPs were separated from unbound plasma components. For polystyrene NPs, separation was achieved by centrifugation at 20000 × g for 30 minutes at 10–15 °C to pellet the NP-protein complexes. For SPION beads, both magnetic separation (using a strong neodymium magnet for 5 minutes) and centrifugation were employed to compare isolation efficacy across methods. To remove loosely bound proteins and non-specific plasma components, the pelleted or magnetically separated NPs were subjected to three rigorous washing steps. Each wash involved carefully resuspending the pellet in 400 μL of cold Sorensen’s phosphate buffer and centrifuging or magnetically separating again. Finally, the extensively washed corona-coated NPs were resuspended in 200 μL of Sorensen’s phosphate buffer for subsequent analysis.

**Dynamic light scattering (DLS).** The hydrodynamic size distribution of the polystyrene NPs was determined using a Zetasizer Nano Series instrument (Malvern Instruments, UK). Measurements were performed at 25°C with a 632 nm Helium-Neon laser.

**Sodium dodecyl sulfate–polyacrylamide gel electrophoresis (SDS-PAGE)**. SDS-PAGE was employed for an initial qualitative assessment of the protein corona composition. Aliquots (20 μL) of the final corona-coated NP suspensions were mixed with 20 μL of 2× Laemmli sample buffer (Bio-Rad), heated at 85°C for 6 minutes, and loaded onto precast 4-12% gradient polyacrylamide gels (NuPAGE, Invitrogen). Following electrophoresis at 120V for 90 minutes, gels were fixed in a solution containing 10% acetic acid and 40% ethanol for 30 minutes, then stained overnight with 50 mL of Coomassie blue stain (Bio-Rad). After destaining with multiple washes of Milli-Q water, the gels were scanned using a gel documentation system for visual analysis of protein banding patterns.

**Cryogenic transmission electron microscopy (Cryo-TEM).** To visualize the morphology of corona-coated NPs and to directly observe EV co-isolation, Cryo-TEM was performed. For imaging, 10-nm BSA-treated gold NPs (Ted Pella, Inc.) were added as fiducial markers to each of the corona-coated NPs at a ratio of 1:4.3. Five microliters of each sample were applied to glow-discharged holey carbon grids (C-Flat R2/2, Protochips, Inc.), blotted to create a thin liquid film (approximately 50 nm thick) using a Vitrobot Mark IV (Thermo Fisher Scientific, Hillsboro, OR, USA), and rapidly frozen in liquid ethane to preserve structural integrity. Images were acquired on a Titan Krios 300 kV Cryo-TEM equipped with a Falcon 2 direct electron detector and phase plate (Thermo Fisher Scientific). The microscope was operated at a nominal magnification of 75,000×, resulting in a pixel size of 1.075 Å, with defocus values ranging from −2.0 to −3.0 μm under low-dose conditions.

**Scanning electron microscopy (SEM).** A 20 µL aliquot of the solution was deposited onto a poly-L-lysine-coated glass coverslip. Following freeze-drying, samples were sputter-coated with a thin layer of platinum to enhance electrical conductivity and minimize charging effects during SEM imaging. Scanning electron microscopy was performed using an FEI Quanta 450 instrument at a low accelerating voltage of 5 kV.

**Proteomics sample preparation.** For comprehensive protein quantification, the isolated NP-protein corona pellets were prepared for mass spectrometry analysis. The pellets were first resuspended in 20 µL of a denaturing buffer containing 0.5 M guanidinium hydrochloride in phosphate-buffered saline (PBS). Proteins were then reduced by adding 2 mM final concentration of dithiothreitol (DTT) and incubated at 37°C for 45 min. Proteins were then alkylated by adding 8 mM final concentration of iodoacetamide (IAA) and incubated for 45 minutes at room temperature in the dark. For digesting the proteins, LysC was added, and the mixture was incubated for 4 hours at 37°C, followed by overnight digestion with 0.1 µg trypsin at 37°C. To separate the resulting peptides from the NPs, the digested samples were then centrifuged at 12,000 × g for 5 minutes at room temperature using Vivaspin 500 filter columns (e.g., with a 10 kDa molecular weight cut-off). The peptide-containing flow-through (bottom fraction) was collected, while the larger NPs and undigested proteins were retained on the filter. The collected peptides were subsequently acidified to pH 2-3 using trifluoroacetic acid (TFA) and purified using C18 cartridges according to the manufacturer's instructions to remove salts and detergents prior to LC-MS/MS analysis.

**LC-MS/MS.**

Dried peptides were resuspended in 0.1% aqueous formic acid, 0.02% DDM (n-Dodecyl-B-D-maltoside) and subjected to LC–MS/MS analysis using a timsTOF Ultra 2 Mass Spectrometer equipped with a CaptiveSpray nano-electrospray ion source (both Bruker) and fitted with a Vanquish Neo (Thermo Fisher Scientific). Peptides were resolved using a RP-HPLC column (100 um × 30 cm) packed in-house with C18 resin (ReproSil Saphir 100 C18, 1.5 um resin; Dr. Maisch GmbH) at a flow rate of 0.4 ul/min and column heater set to 60°C. The following gradient was used for peptide separation: from 2% B to 25% B over 25 min to 35% B over 5 min to 95% B over 0.5 min followed by 4 min at 95% B to 2% B over 0.5 min followed by 0.5 min at 2% B. Buffer A was 0.1% formic acid in water and buffer B was 80% acetonitrile, 0.1% formic acid in water.

The mass spectrometer was operated in dia-PASEF mode with a cycle time estimate of 0.95 s. MS1 and MS2 scans were acquired over a mass range from 100 to 1700 m/z. A method with 8 dia-PASEF scans separated into 3 ion mobility windows per scan covering a 400-1000 m/z range with 25 Da windows and an ion mobility range from 0.64 to 1.37 Vs cm^2^ was used. Ion Charge Control (ICC 2.0) was enabled. Accumulation and ramp time were set to 100 ms, capillary voltage was set to 1600V, dry gas was set to 3 l/min and dry temperature was set to 200 °C. The collision energy was ramped linearly as a function of ion mobility from 59 eV at 1/K0 = 1.6 V s cm^-2^ to 20 eV at 1/K0 = 0.6 V s cm^-2^.

The acquired files were searched using the Spectronaut (Biognosys v19.0) directDIA workflow against a Homo sapiens database (consisting of 20360 protein sequences downloaded from Uniprot on 20220222) and 392 commonly observed contaminants using default settings except Cross-Run Normalization was disabled. Quantitative data was exported from Spectronaut using the Pivot Report export function.

**Data analysis**

Raw proteomic datasets were received with three replicates per sample. All data processing and analysis were performed using custom Python scripts. All graphing and statistical analyses were performed using GraphPad Prism (version 10.6.1). UniProt was used as the reference database for protein identifiers, annotations, and functional classification. Prior to quantitative analysis, contaminant proteins were removed based on annotation, excluding entries containing the prefix “Con–” in the protein description. Missing values were subsequently imputed at the replicate level: when a single replicate value was missing, it was imputed using the average of the remaining two replicates; proteins missing values in two or more replicates were excluded from analysis for that sample. Protein abundances were then normalized by total protein intensity within each replicate and expressed as relative abundance percentages. These normalized values were used for all downstream analyses to enable compositional comparisons across experimental conditions. Z-score normalization was performed on replicate-averaged data by centering and scaling the abundance of each protein across all conditions.

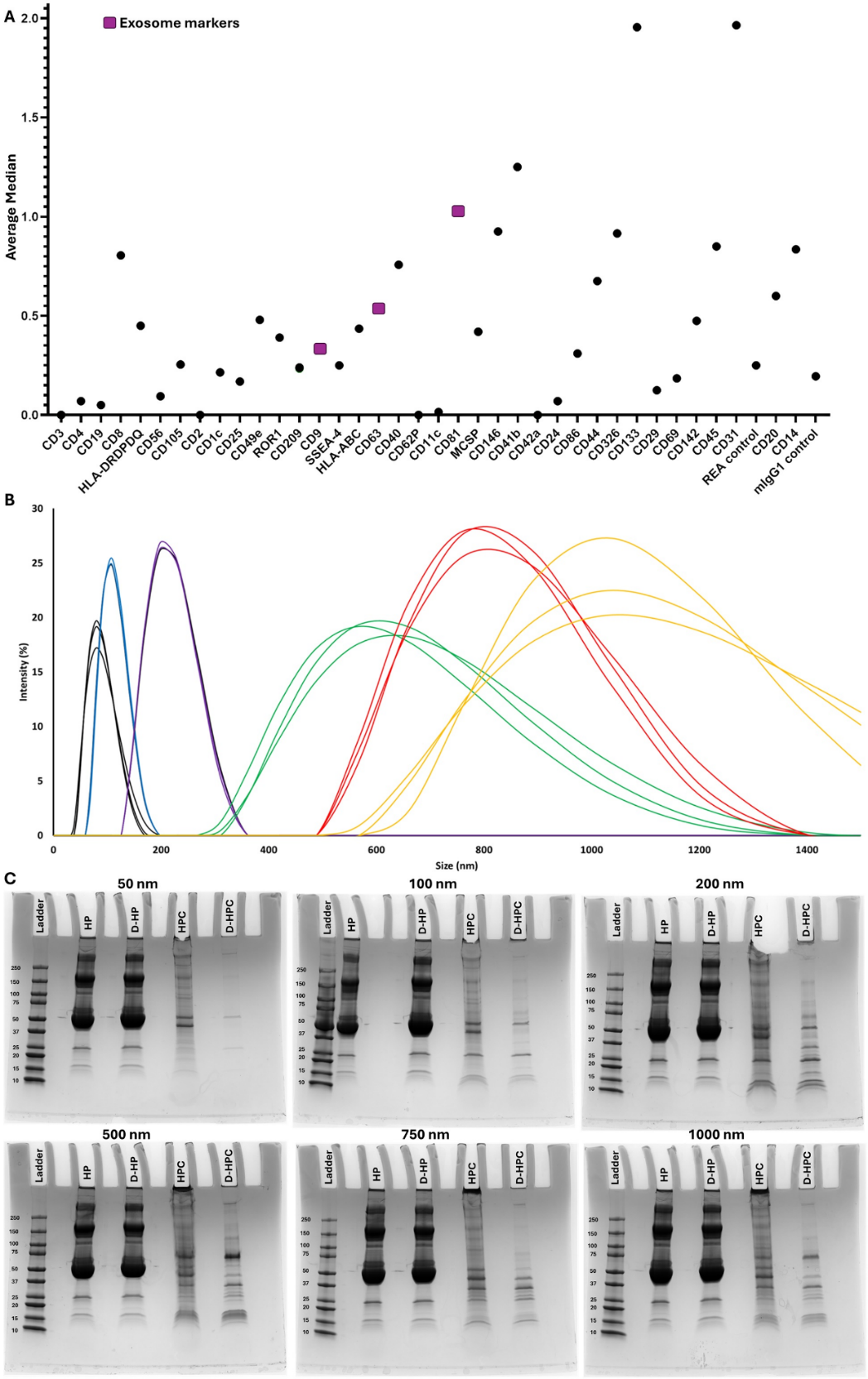

**Figure S1. Characterization of plasma conditions and NP corona formation.** (A) Characterization of the 100,000 × g plasma pellet. MACSPlex immunoaffinity analysis of the resuspended pellet confirms enrichment of material bearing 37 EV-associated surface epitopes, including strong signals for canonical tetraspanin exosome markers CD9, CD63, and CD81. (B) Size distribution analysis of polystyrene NPs of various sizes via DLS. (C) SDS-PAGE profiles of proteins recovered from corona-coated NPs incubated in regular/neat versus EV-depleted plasma, revealing distinct molecular weight distributions, with several bands present in neat plasma being absent or diminished in the EV-depleted condition. HP: Human Plasma; D-HP: Depleted Human Plasma; HPC: Human Plasma Corona; D-HPC: Depleted Human Plasma Corona.

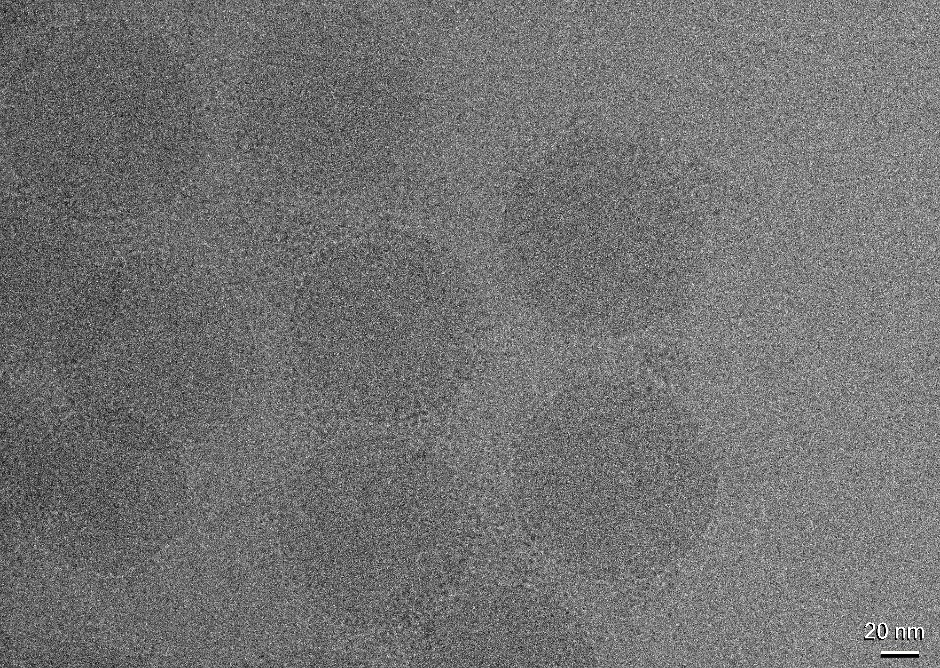

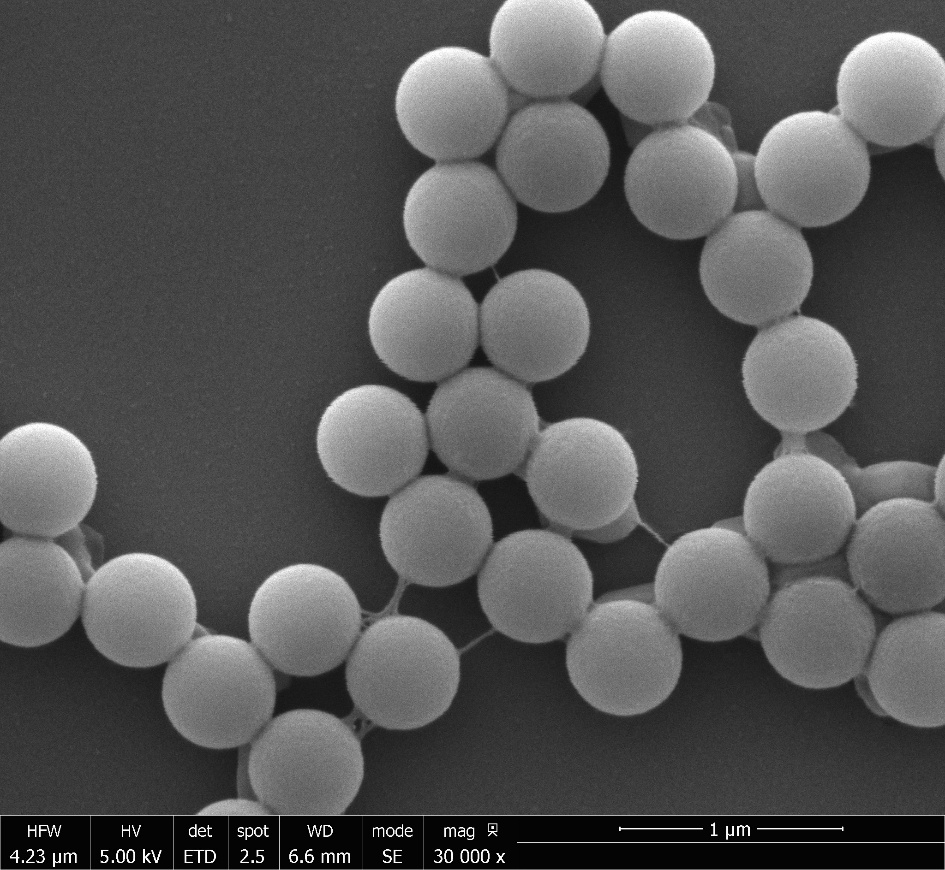

**Figure S2. Imaging of the protein corona after incubation with EV-depleted plasma.** Representative imaging of corona-coated polystyrene NPs after incubation with EV-depleted plasma using cryo-TEM (top panel) and SEM (bottom panel).

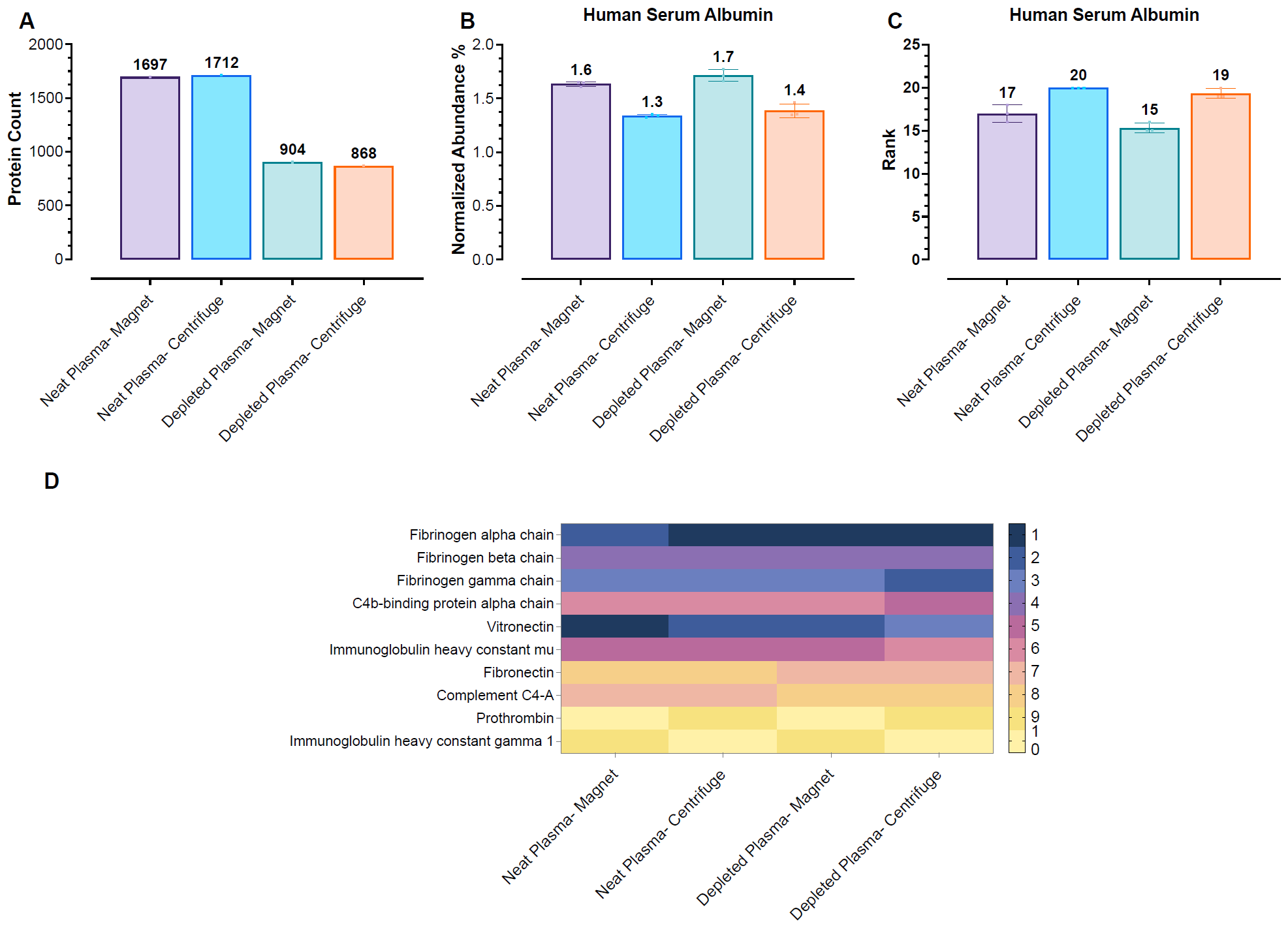

**Figure S3. Impact of EV depletion on SPION beads’ protein corona analysis across different isolation methods.** (A) Total count of unique proteins identified via LC-MS/MS on SPION beads. Coronas were isolated by either magnetic separation or centrifugation from neat or EV-depleted plasma. (B) Normalized spectral abundance of albumin on magnetic beads. (C) The rank of albumin within the total identified SPION beads corona proteins. (D) Heatmap of the top 10 most abundant proteins identified in each SPION beads (Rank 1 = Dark Blue; Rank 10 = Yellow). The top proteins are primarily highly abundant plasma proteins involved in coagulation, complement, and immunity (e.g., Fibrinogen, C4b-binding protein, Immunoglobulins, Fibronectin).

**Table S1. The name and ID of FDA-approved and diagnostic biomarkers**

| UniProt_ID | Protein_name |
| --- | --- |
| P00338 | L-lactate dehydrogenase A chain |
| P00450 | Ceruloplasmin |
| P00488 | Coagulation factor XIII A chain |
| P00738 | Haptoglobin |
| P00740 | Coagulation factor IX |
| P00742 | Coagulation factor X |
| P00747 | Plasminogen |
| P00749 | Urokinase-type plasminogen activator |
| P00750 | Tissue-type plasminogen activator |
| P00797 | Renin |
| P01008 | Antithrombin-III |
| P01009 | Alpha-1-antitrypsin |
| P01023 | Alpha-2-macroglobulin |
| P01024 | Complement C3 |
| P01031 | Complement C5 |
| P01034 | Cystatin-C |
| P01215 | Glycoprotein hormones alpha chain |
| P01222 | Thyrotropin subunit beta |
| P01229 | Lutropin subunit beta |
| P01236 | Prolactin |
| P01241 | Somatotropin |
| P01258 | Calcitonin |
| P01266 | Thyroglobulin |
| P01270 | Parathyroid hormone |
| P01344 | Insulin-like growth factor 2 |
| P01588 | Erythropoietin |
| P01589 | Interleukin-2 receptor subunit alpha |
| P02144 | Myoglobin |
| P02647 | Apolipoprotein A-I |
| P02671 | Fibrinogen alpha chain |
| P02675 | Fibrinogen beta chain |
| P02741 | C-reactive protein |
| P02745 | Complement C1q subcomponent subunit A |
| P02746 | Complement C1q subcomponent subunit B |
| P02747 | Complement C1q subcomponent subunit C |
| P02751 | Fibronectin |
| P02753 | Retinol-binding protein 4 |
| P02763 | Alpha-1-acid glycoprotein 1 |
| P02765 | Alpha-2-HS-glycoprotein |
| P02766 | Transthyretin |
| P02768 | Albumin |
| P02771 | Alpha-fetoprotein |
| P02775 | Platelet basic protein |
| P02786 | Transferrin receptor protein 1 |
| P02787 | Serotransferrin |
| P02790 | Hemopexin |
| P02792 | Ferritin light chain |
| P02794 | Ferritin heavy chain |
| P02818 | Osteocalcin |
| P04070 | Vitamin K-dependent protein C |
| P04075 | Fructose-bisphosphate aldolase A |
| P04114 | Apolipoprotein B-100 |
| P04275 | von Willebrand factor |
| P04278 | Sex hormone-binding globulin |
| P04626 | Receptor tyrosine-protein kinase erbB-2 |
| P04746 | Pancreatic alpha-amylase |
| P05062 | Fructose-bisphosphate aldolase B |
| P05111 | Inhibin alpha chain |
| P05121 | Plasminogen activator inhibitor 1 |
| P05155 | Plasma protease C1 inhibitor |
| P05160 | Coagulation factor XIII B chain |
| P05164 | Myeloperoxidase |
| P05186 | Alkaline phosphatase, tissue-nonspecific isozyme |
| P06276 | Cholinesterase |
| P06732 | Creatine kinase M-type |
| P06744 | Glucose-6-phosphate isomerase |
| P07195 | L-lactate dehydrogenase B chain |
| P07225 | Vitamin K-dependent protein S |
| P07288 | Prostate-specific antigen |
| P07477 | Serine protease 1 |
| P08519 | Apolipoprotein(a) |
| P08697 | Alpha-2-antiplasmin |
| P08833 | Insulin-like growth factor-binding protein 1 |
| P0C0L4 | Complement C4-A |
| P12277 | Creatine kinase B-type |
| P12821 | Angiotensin-converting enzyme |
| P13686 | Tartrate-resistant acid phosphatase type 5 |
| P14618 | Pyruvate kinase PKM |
| P15309 | Prostatic acid phosphatase |
| P15941 | Mucin-1 |
| P16233 | Pancreatic triacylglycerol lipase |
| P16860 | Natriuretic peptides B |
| P17174 | Aspartate aminotransferase, cytoplasmic |
| P17936 | Insulin-like growth factor-binding protein 3 |
| P19429 | Troponin I, cardiac muscle |
| P19440 | Glutathione hydrolase 1 proenzyme |
| P19652 | Alpha-1-acid glycoprotein 2 |
| P24298 | Alanine aminotransferase 1 |
| P43251 | Biotinidase |
| P45379 | Troponin T, cardiac muscle |
| P61626 | Lysozyme C |
| P61769 | Beta-2-microglobulin |
| Q00796 | Sorbitol dehydrogenase |
| Q14508 | WAP four-disulfide core domain protein 2 |
| Q8WXI7 | Mucin-16 |
| O75874 | Isocitrate dehydrogenase [NADP] cytoplasmic |
| P00390 | Glutathione reductase, mitochondrial |
| P00451 | Coagulation factor VIII |
| P00505 | Aspartate aminotransferase, mitochondrial |
| P00734 | Prothrombin |
| P00748 | Coagulation factor XII |
| P01019 | Angiotensinogen |
| P01116 | GTPase KRas |
| P01189 | Pro-opiomelanocortin |
| P01350 | Gastrin |
| P01730 | T-cell surface glycoprotein CD4 |
| P01889 | HLA class I histocompatibility antigen, B alpha chain |
| P01903 | HLA class II histocompatibility antigen, DR alpha chain |
| P01909 | HLA class II histocompatibility antigen, DQ alpha 1 chain |
| P01911 | HLA class II histocompatibility antigen, DRB1 beta chain |
| P01920 | HLA class II histocompatibility antigen, DQ beta 1 chain |
| P02649 | Apolipoprotein E |
| P02749 | Beta-2-glycoprotein 1 |
| P02760 | Protein AMBP |
| P02776 | Platelet factor 4 |
| P02788 | Lactotransferrin |
| P03951 | Coagulation factor XI |
| P03952 | Plasma kallikrein |
| P04040 | Catalase |
| P04062 | Lysosomal acid glucosylceramidase |
| P04439 | HLA class I histocompatibility antigen, A alpha chain |
| P05019 | Insulin-like growth factor 1 |
| P05106 | Integrin beta-3 |
| P05109 | Protein S100-A8 |
| P05455 | Lupus La protein |
| P05556 | Integrin beta-1 |
| P06280 | Alpha-galactosidase A |
| P06702 | Protein S100-A9 |
| P06881 | Calcitonin gene-related peptide 1 |
| P07359 | Platelet glycoprotein Ib alpha chain |
| P07478 | Trypsin-2 |
| P07864 | L-lactate dehydrogenase C chain |
| P07902 | Galactose-1-phosphate uridylyltransferase |
| P08236 | Beta-glucuronidase |
| P08514 | Integrin alpha-IIb |
| P08575 | Receptor-type tyrosine-protein phosphatase C |
| P08709 | Coagulation factor VII |
| P08727 | Keratin, type I cytoskeletal 19 |
| P09455 | Retinol-binding protein 1 |
| P09619 | Platelet-derived growth factor receptor beta |
| P09923 | Intestinal-type alkaline phosphatase |
| P09936 | Ubiquitin carboxyl-terminal hydrolase isozyme L1 |
| P09972 | Fructose-bisphosphate aldolase C |
| P0C0L5 | Complement C4-B |
| P10253 | Lysosomal alpha-glucosidase |
| P10635 | Cytochrome P450 2D6 |
| P10696 | Alkaline phosphatase, germ cell type |
| P10745 | Retinol-binding protein 3 |
| P11117 | Lysosomal acid phosphatase |
| P11388 | DNA topoisomerase 2-alpha |
| P11413 | Glucose-6-phosphate 1-dehydrogenase |
| P12259 | Coagulation factor V |
| P12532 | Creatine kinase U-type, mitochondrial |
| P13224 | Platelet glycoprotein Ib beta chain |
| P13762 | HLA class II histocompatibility antigen, DR beta 4 chain |
| P14136 | Glial fibrillary acidic protein |
| P14770 | Platelet glycoprotein IX |
| P14780 | Matrix metalloproteinase-9 |
| P14784 | Interleukin-2 receptor subunit beta |
| P15056 | Serine/threonine-protein kinase B-raf |
| P16671 | Platelet glycoprotein 4 |
| P17301 | Integrin alpha-2 |
| P17540 | Creatine kinase S-type, mitochondrial |
| P17931 | Galectin-3 |
| P18146 | Early growth response protein 1 |
| P19961 | Alpha-amylase 2B |
| P20061 | Transcobalamin-1 |
| P20815 | Cytochrome P450 3A5 |
| P21589 | 5'-nucleotidase |
| P21980 | Protein-glutamine gamma-glutamyltransferase 2 |
| P22303 | Acetylcholinesterase |
| P22309 | UDP-glucuronosyltransferase 1A1 |
| P24158 | Myeloblastin |
| P24666 | Low molecular weight phosphotyrosine protein phosphatase |
| P28838 | Cytosol aminopeptidase |
| P28908 | Tumor necrosis factor receptor superfamily member 8 |
| P30613 | Pyruvate kinase PKLR |
| P31785 | Cytokine receptor common subunit gamma |
| P35030 | Trypsin-3 |
| P35475 | Alpha-L-iduronidase |
| P40197 | Platelet glycoprotein V |
| P40692 | DNA mismatch repair protein Mlh1 |
| P42898 | Methylenetetrahydrofolate reductase (NADPH) |
| P43246 | DNA mismatch repair protein Msh2 |
| P48735 | Isocitrate dehydrogenase [NADP], mitochondrial |
| P50120 | Retinol-binding protein 2 |
| P51580 | Thiopurine S-methyltransferase |
| P52701 | DNA mismatch repair protein Msh6 |
| P54278 | Mismatch repair endonuclease PMS2 |
| P54803 | Galactocerebrosidase |
| P68871 | Hemoglobin subunit beta |
| P69905 | Hemoglobin subunit alpha |
| P79483 | HLA class II histocompatibility antigen, DR beta 3 chain |
| P82980 | Retinol-binding protein 5 |
| Q01638 | Interleukin-1 receptor-like 1 |
| Q02161 | Blood group Rh(D) polypeptide |
| Q12882 | Dihydropyrimidine dehydrogenase [NADP(+)] |
| Q13093 | Platelet-activating factor acetylhydrolase |
| Q14980 | Nuclear mitotic apparatus protein 1 |
| Q16671 | Anti-Muellerian hormone type-2 receptor |
| Q16849 | Receptor-type tyrosine-protein phosphatase-like N |
| Q30154 | HLA class II histocompatibility antigen, DR beta 5 chain |
| Q5S007 | Leucine-rich repeat serine/threonine-protein kinase 2 |
| Q6ZMR3 | L-lactate dehydrogenase A-like 6A |
| Q6ZNF0 | Acid phosphatase type 7 |
| Q8N4E7 | Ferritin, mitochondrial |
| Q8NHS2 | Putative aspartate aminotransferase, cytoplasmic 2 |
| Q96R05 | Retinoid-binding protein 7 |
| Q9BQB6 | Vitamin K epoxide reductase complex subunit 1 |
| Q9BYZ2 | L-lactate dehydrogenase A-like 6B |
| Q9BZG2 | Testicular acid phosphatase |
| Q9BZJ3 | Tryptase delta |
| Q9NPH0 | Lysophosphatidic acid phosphatase type 6 |
| Q9NZQ7 | Programmed cell death 1 ligand 1 |
| Q9UM73 | ALK tyrosine kinase receptor |
| Q9UP52 | Transferrin receptor protein 2 |
| Q9Y6L6 | Solute carrier organic anion transporter family member 1B1 |
| P00751 | Complement factor B |
| P01343 | Insulin-like growth factor 1 |
| P02679 | Fibrinogen gamma chain |
| P04054 | Phospholipase A2 |
| P05543 | Thyroxine-binding globulin |
| P06731 | Cell adhesion molecule CEACAM5 |
| P0DOX7 | Immunoglobulin kappa light chain |
| P0DOX8 | Immunoglobulin lambda-1 light chain |
| Q13421 | Mesothelin |
| Q5EK51 | Lactotransferrin |

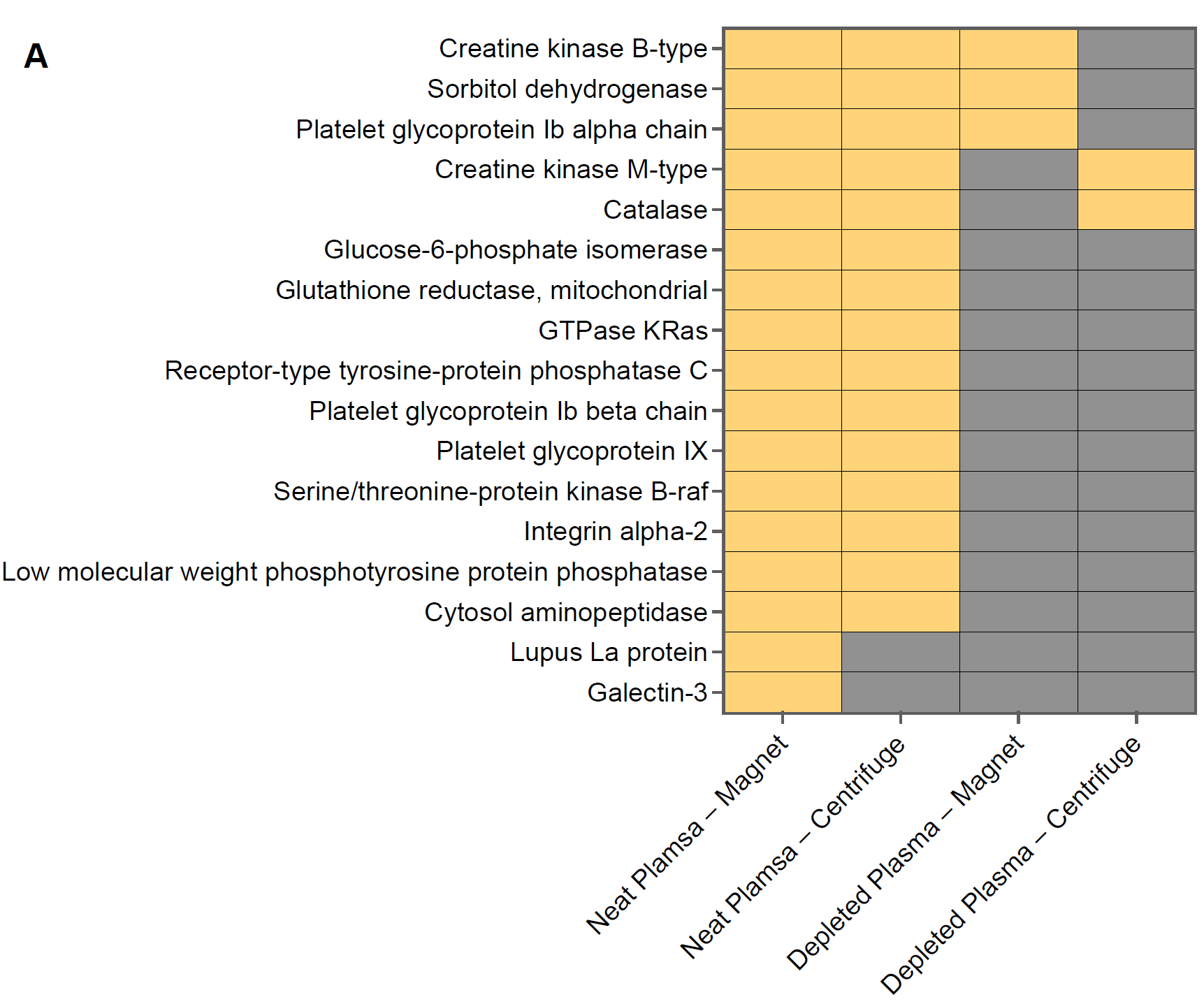

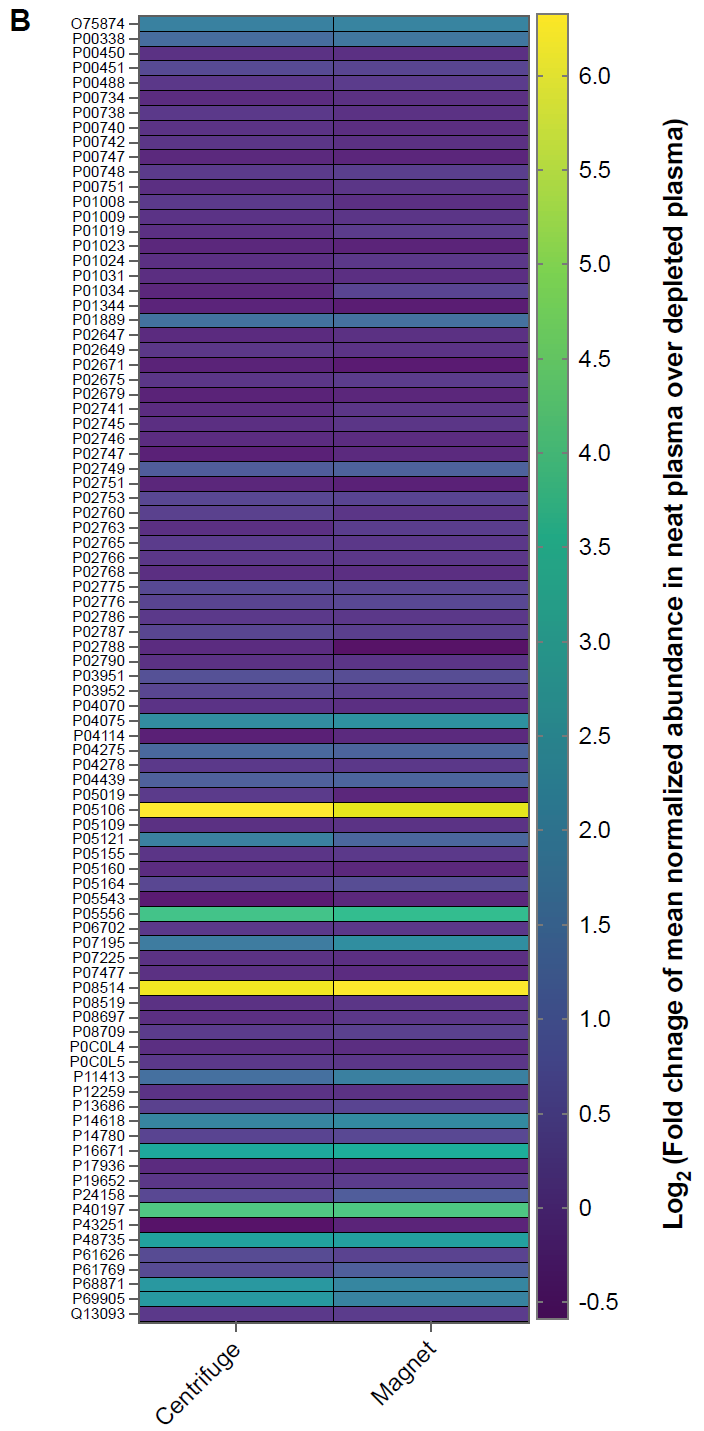

**Figure S4. EVs’ influence on detection of FDA-approved biomarkers in protein coronas of SPION beads.** (A) Heatmap illustrating the detection (yellow) or absence (gray) of 4 FDA-approved and diagnostic plasma biomarkers in the protein corona of SPION beads. The horizontal axis represents different separation techniques (magnetic separation and centrifugation) and plasma conditions (neat plasma vs. depleted plasma). (B) Heatmap showing the Log_2_ of the fold change of the normalized abundance in neat plasma over EV-depleted plasma for FDA-approved and clinical plasma biomarkers commonly detected across nanoparticle protein coronas. For each nanoparticle size, replicate measurements were averaged prior to calculation. Positive values indicate higher enrichment in neat plasma, whereas negative values indicate higher enrichment in EV-depleted plasma.

**References**

1.	H. Tang, J. Wang, M. Mahmoudi, Improving accuracy and reproducibility of mass spectrometry characterization of protein coronas on nanoparticles. *Nature Protocols* **20**, 3057–3063 (2025).

2.	S. Sheibani *et al.*, Nanoscale characterization of the biomolecular corona by cryo-electron microscopy, cryo-electron tomography, and image simulation. *Nature Communications* **12**, 573 (2021).
